# Two neuropeptide signalings regulate post-mating refusal behavior and reproductive system in male crickets

**DOI:** 10.1101/2024.12.20.629589

**Authors:** Zhen Zhu, Shinji Nagata

## Abstract

After mating, insects always perform mating refusal behavior, termed post-mating refractoriness, due to physiological restrictions. Male crickets, *Gryllus bimaculatus*, characteristically exhibit 1-hour post-mating refractory stage, controlled by terminal abdominal ganglion. The molecular mechanisms underlying the male-specific precisely timed refractory stage remain elucidated. Here we show that among 28 neuropeptide precursors expressed in the terminal abdominal ganglion, *DH31*, *myosuppressin*, *allatotropin*, and *sNPF* exhibited male-specific expression based on RT-qPCR and *in situ* hybridization. However, RNA interference experiments showed that only knockdown of *allatotropin* and *sNPF* changed the duration of the refractoriness. Furthermore, *allatotropin* and *sNPF* knockdown influenced functions of male reproductive system by inhibiting seminal fluid secretion from male accessory gland and decreasing sperm storage in seminal vesicles, respectively. Knockdown of their receptors caused similar phenotypes. In conclusion, this study demonstrated the regulation of post-mating refusal behavior and reproductive system by Allatotropin and sNPF signalings in male crickets.

## Introduction

In polygamous insects, multiple matings benefit the fecundity and fertility of the population ^1^. Females with experience of multiple matings are guarded by more mating partners and protected against male infertility, and replenish the depleted sperm stores ^2,3^. Additionally, multiple matings contribute to male competition for fertilization success ^4^. Despite these advantages, the energy- consumption of mating restricts continuous mating behavior in most cases. Therefore, immediately after mating, insects enter a refractory stage (RS) during which they perform mating refusal behavior and replenish the energy and gametes consumed in the previous mating. The duration of the post-mating refractoriness limits the mating frequency. Extensive research has focused on the regulatory mechanisms underlying post-mating refractoriness in females. Female insects typically receive special substances from males, such as sex peptide from the male fruit fly *Drosophila melanogaster* and a male-specifically modified steroid, 3-dehydro-20- hydroxyecdysone from the male mosquito, *Anopheles gambiae*, during copulation ^5,6^. These male-derived substances activate downstream singaling in females to reduce receptivity to remating, leading to post-mating refractoriness. In contrast, much less attention has been given to understanding the mechanisms underlying post-mating refractoriness in male insects.

Crickets, belonging to the Grylloidea insect superfamily, serve as excellent models for studying male post-mating refractoriness due to their identifiable reproductive cycle which is composed of mating stage and post-mating RS. The mating stage initiates with a calling or courtship song and ends with spermatophore extrusion, while the RS begins with spermatophore extrusion and extends to the onset of next calling or courtship songs ^7^ (Fig. 1). Crucially, different from the female refractoriness, whose duration changes with the accumulation of male-derived substances ^8^, the RS of male crickets is precisely timed ^9^. This stage can be further compartmentalized into RS1, which is from the extrusion of the old spermatophore to the protrusion of a new immature spermatophore, and RS2, which is from spermatophore protrusion to the onset of calling or courtship songs (Fig. 1). In the case of the two-spotted cricket, *Gryllus bimaculatus*, although the duration of RS1 is susceptible to internal and external influences, it typically lasts for approximately 6 minutes under normal conditions in the presence of females ^10^. The duration of RS2 in *G. bimaculatus* is constantly 1 hour ^11^. The clock timer governing the RS2 has been identified in the terminal abdominal ganglion (TAG), located at the end of the central nervous system in crickets ^12^. However, the molecular mechanisms underlying the precisely timed post-mating refractoriness remain largely unknown.

**Figure 1.**
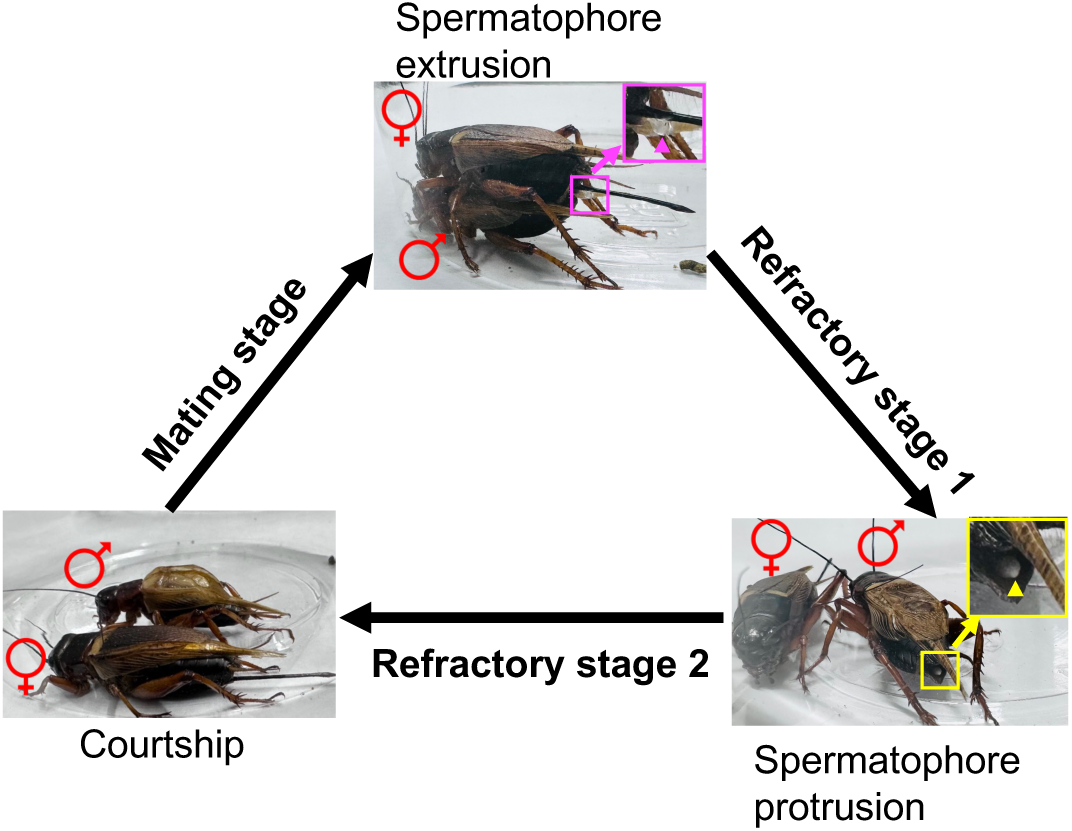
The male reproductive cycle of *G. bimaculatus*. Mating stage is from calling or courtship observed by wing vibration to spermatophore extrusion of males. Magenta rectangle region is enlarged. Magenta triangle points toward the spermatophore that is extruded from the male and attached to the external genital of the mated female. Refractory stage is further divided into stage 1 (RS1) and stage 2 (RS2). RS1 is from spermatophore extrusion to the protrusion of a new immature spermatophore which is enlarged and pointed by yellow triangle. RS2 is from spermatophore protrusion to the onset of calling or courtship.

Neuropeptides are the largest group of critical signaling molecules that orchestrate various physiological and behavioral activities, including reproduction in animals ^13^. To date, few descriptions about the regulation of reproductive behaviors in male insects by neuropeptides have been reported. We here sought to decipher whether TAG-derived neuropeptides were involved in regulating the precisely timed male post-mating refractoriness and explore their functions.

In the present study, we identified 28 neuropeptide precursors expressed in the TAG. Two neuropeptides, *allatotropin* and *snpf*, contributed to the precisely timed male post-mating refractoriness. Additionally, the two neuropeptides regulated the physiological activities in the male reproductive system that are essential for spermatophore preparation. Specifically, knockdown of *allatotropin* signaling suppressed secretion of seminal fluid from the male accessory gland, while knockdown of *snpf* signaling inhibited sperm storage in the seminal vesicles. In summary, our findings provide new insights into the mechanisms underlying the male post-mating refractoriness and spermatophore preparation in the male reproductive system.

## Results

### *Myosuppressin, AT, DH31,* and *sNPF* exhibit male-specific expression in the TAG

To investigate the regulation of male-specific, precisely timed post-mating refractoriness by neuropeptides in crickets, we first explored the neuropeptides with sexually different expression. In the TAGs from male adults on the 4th day after eclosion with sexual maturation, 28 neuropeptide precursors were identified by RNA sequencing analysis, and subsequent RT-qPCR unveiled that 4 of 28 neuropeptide precursors exhibited sexually different expression in the TAGs (Fig. 2); *Adipokinetic hormone/corazonin-related peptide* (*ACP*) showed lower expression in the male TAG than in the female TAG, whereas *myosuppressin*, *allatotropin* (*AT*), and *short neuropeptide F* (*sNPF*) exhibited significantly higher transcriptional levels in the male TAG compared to the female TAG (Fig. 2). The temporal distribution analyses showed that the sexual differences in the transcriptional levels of *myosuppressin*, *AT*, and *sNPF* precursors were observed from the penultimate instar, when the expression of the three precursors increased in the male TAG but did not alter in the female TAG, suggesting their involvement in regulating male-specific events from this developmental stage. Moreover, *AT* and *sNPF* displayed an additional increase in expression at the adult stage, implying potential roles in male adult behaviors and physiological activities (Supplementary Fig. 1 and Fig. 3a-3c).

**Figure 2.**
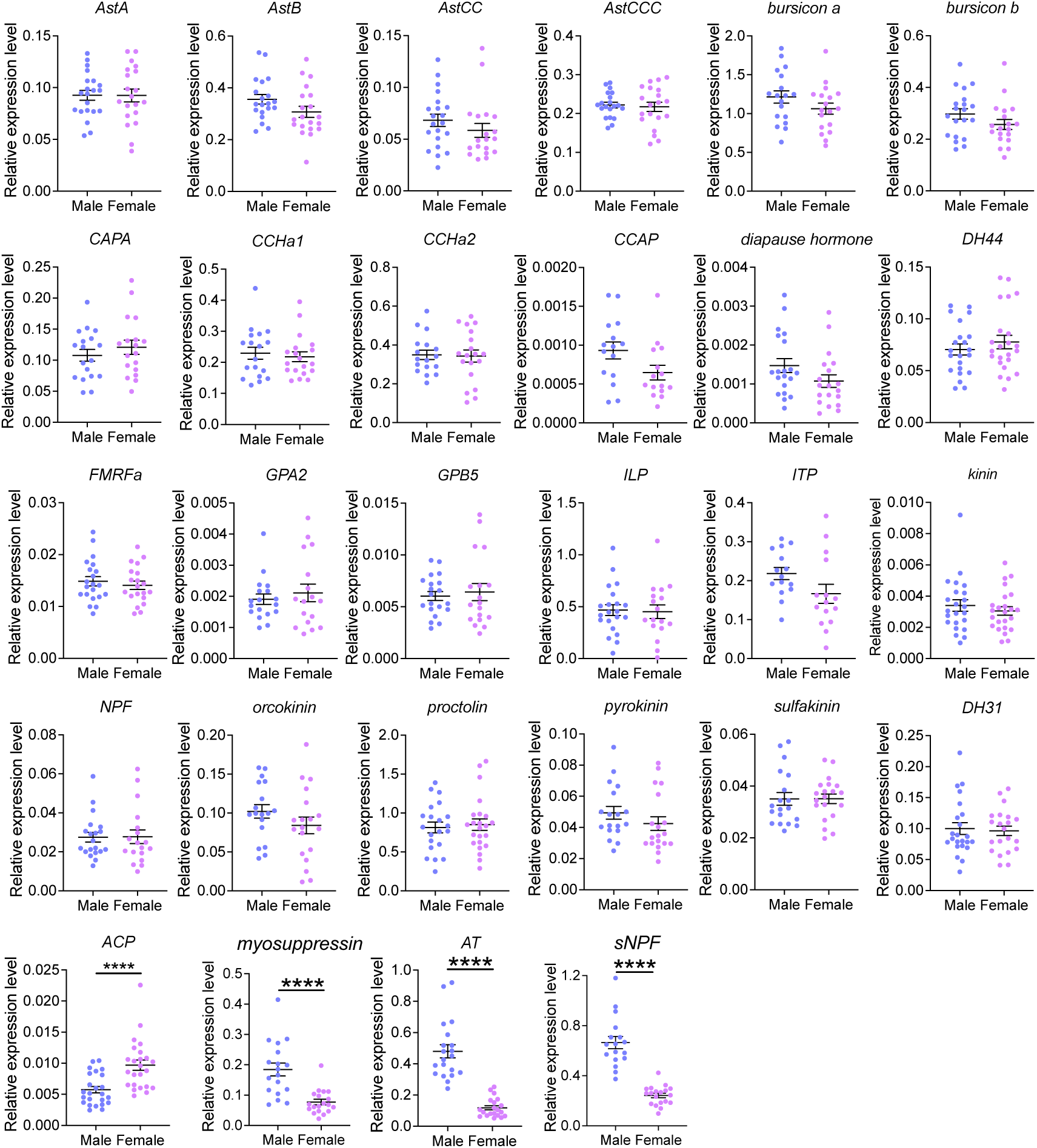
Transcriptional levels of neuropeptide precursors in male and female TAGs. Values are shown as mean ± SEM, n=15-24, unpaired *t*-test, \*\*\*\**P*<0.0001. *AstA, allatostatin A. AstB, allatostatin B. AstCC, allatostatin CC. AstCCC, allatostatin CCC. CAPA, capability. CCHa1, CCHamide1. CCHa2, CCHamide2. CCAP, crustacean cardioactive peptide. DH44, diuretic hormone 44. GPA2, glycoprotein alpha 2. GPB5, glycoprotein beta 5. ILP, insulin-like peptide. ITP, ion transport peptide. NPF, neuropeptide F. DH31, diuretic hormone 31. ACP, adipokinetic hormone/corazonin-related peptide. AT, allatotropin. sNPF, short neuropeptide F*.

**Figure 3.**
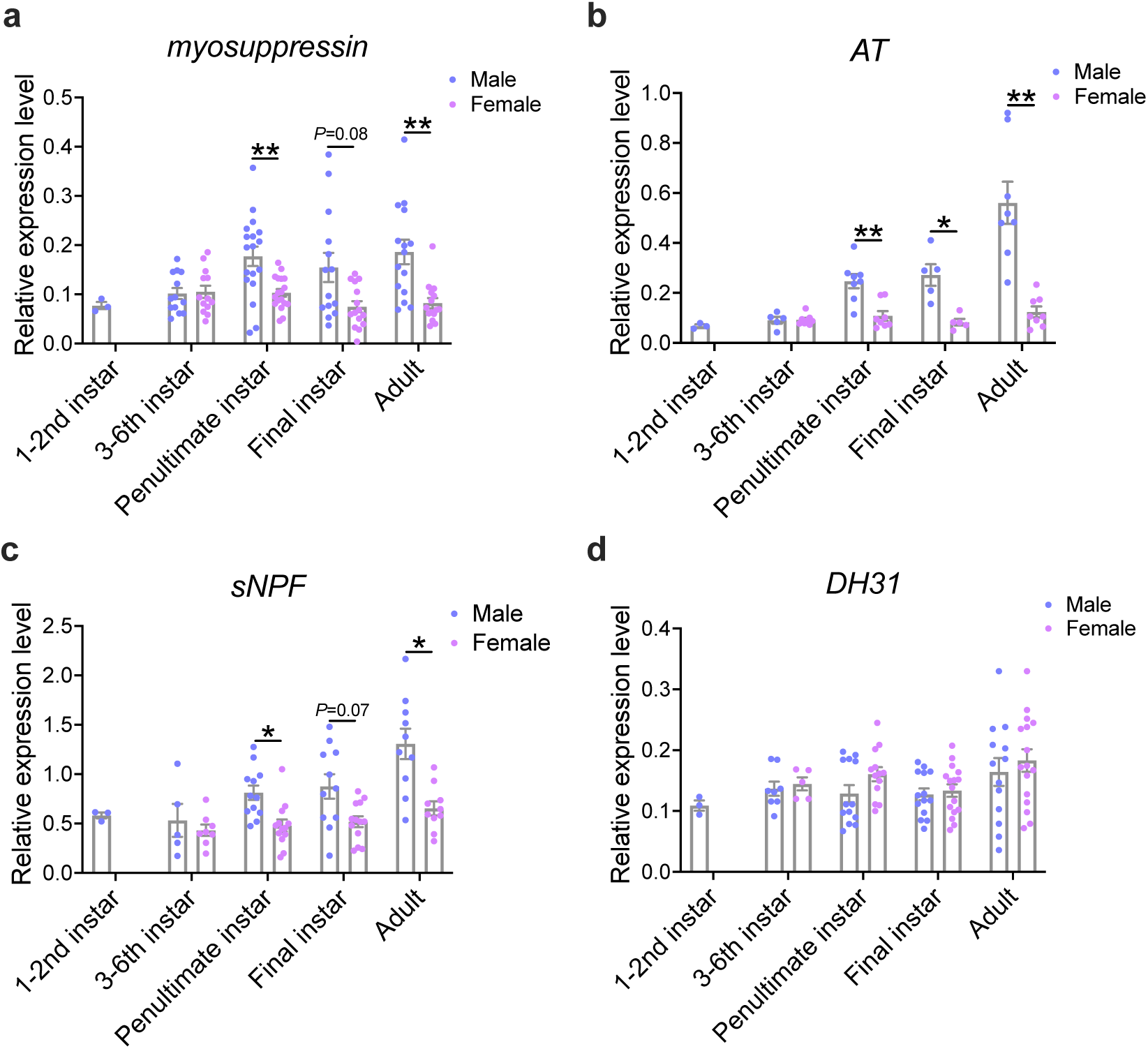
Transcriptional levels of *myosuppressin* (a), *AT* (b), *sNPF* (c), and *DH31* (d) in the TAGs of male and female crickets at different growth stages. Values are shown as mean ± SEM, n=3–18, Sidak’s test, \**P*<0.05, \*\**P*<0.01.

Subsequently, whole-mount *in situ* hybridization revealed the locations of the positive cells expressing the neuropeptide precursors. Among 17 neuropeptide precursors with successfully detected signals, *myosuppressin-*, *AT-*, *sNPF-*, and *diuretic hormone 31* (*DH31*)*-*expressing cells in the TAG exhibited sexual difference (Fig. 4). The positive cells of *myosuppressin*, *AT*, and *sNPF* in the male TAG were noticeably more than those in the female TAG, consistent with the higher transcriptional levels in the male TAG as showed in the RT-qPCR results (Fig. 2). The expressing cells of *myosuppressin*, *AT*, *sNPF*, and *DH31* were similarly distributed in the dorsal midline of the male TAG, forming two nervous cell clusters at the posterior median and bottom median regions. In contrast, no comparable signals were observed in these regions of the female TAG (Fig. 4). In fact, *DH31-*expressing cells were predominantly distributed in the ventral and bilateral regions but not dorsal region, which was observed both in the male and female TAGs (Fig. 4). The difference in the expression in the dorsal TAG might be too small to be detected by RT-qPCR, explaining the comparable *DH31* transcriptional levels in the male and female TAGs at all developmental stages (Fig. 2, 3d, and 4). These findings collectively suggest that the four male-specifically expressing neuropeptides possess the potential of the pivotal roles in male-specific behaviors and physiological activities.

**Figure 4.**
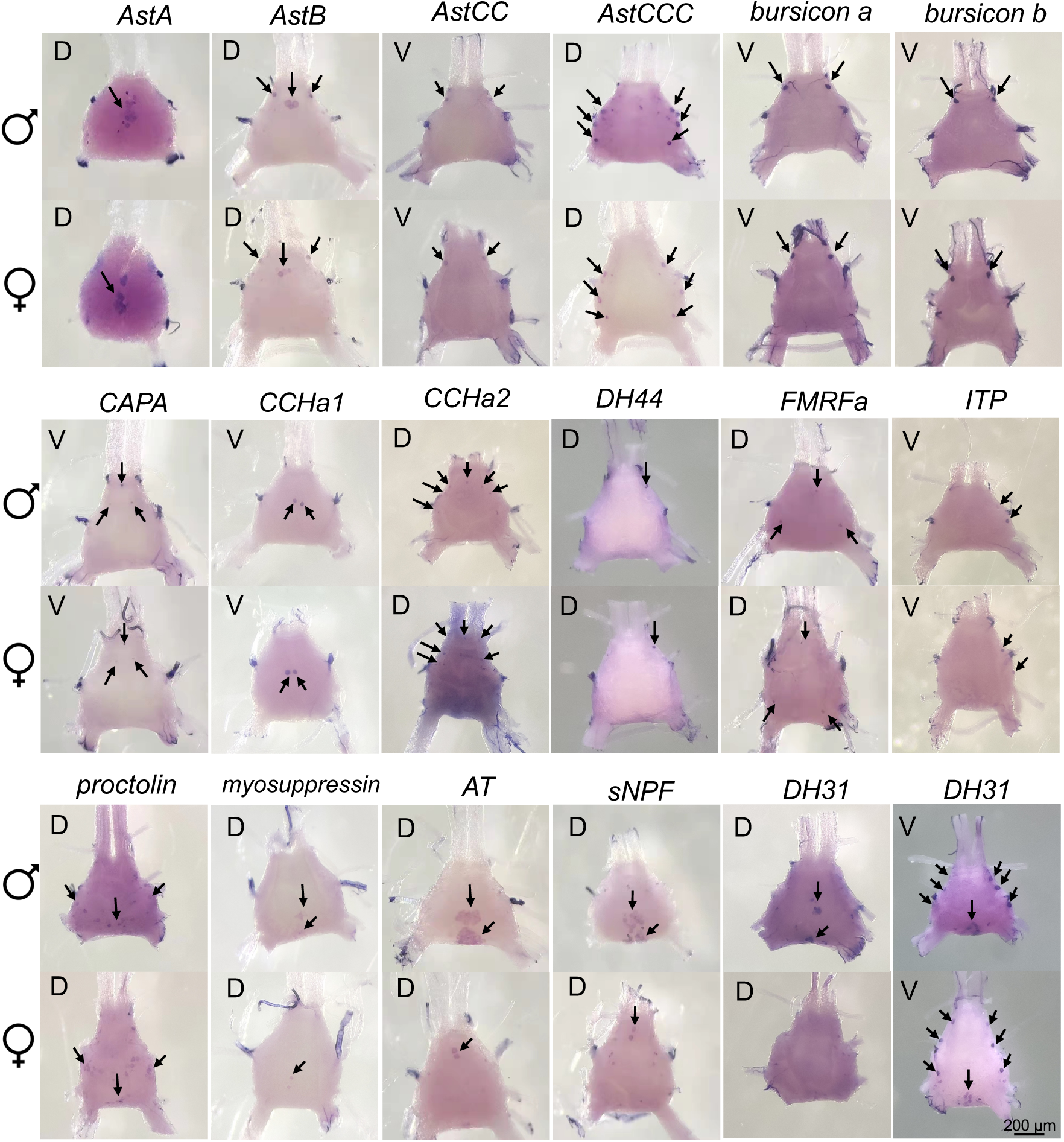
Distributions of neuropeptide*-*expressing cells in male and female TAGs observed by whole-mount *in situ* hybridization. Arrow points to the positive cell. D, dorsal side. V, ventral side.

### AT and sNPF signalings regulate the duration of male post-mating refractoriness

Knockdown experiments using RNA interference (RNAi) were conducted to investigate the contributions of *myosuppressin*, *AT*, *sNPF*, and *DH31* to post-mating refractoriness in male crickets. RNAi was efficient for loss-of-function analyses of these four genes (Supplementary Fig. 2a-2d). The number of matings completed in 12 h and the duration of each post-mating RS were recorded. During 12 h of observation period, *DH31* knockdown males exhibited courtship behavior but failed to transfer spermatophores to females, probably because the male crickets with female mounting were unable to twist their body to hook the epiphallus to the subgenital plate of females for the spermatophore transfer ^14^. These males with failed mating continued courting and did not enter post-mating RS. Consequently, the effects of DH31 on the RS duration could not be evaluated. *Myosuppressin* knockdown males averagely experienced 10.4 matings in 12 h experiment, which was comparable to that of the control males with 11.2 matings (Fig. 5a). Little difference in the duration of RSs was observed between *myosuppressin* knockdown crickets and the control (Fig. 5b). Conversely, *AT* and *sNPF* knockdown males experienced 9.4 and 8.7 matings, respectively, significantly fewer than the control males (Fig. 5a), and exhibited the extended RSs (Fig. 5c and 5d). These results demonstrated that *AT* and *sNPF* knockdown males mated at lower frequency. Like the effects of *AT* and *sNPF* knockdown, the efficient RNAi targeting *AT receptor* (*ATR*) and *sNPF receptor* (*sNPFR*) also decreased the number of matings and extended the RS duration (Supplementary Fig. 2e and 2f; Fig. 5e-5g). As both AT and sNPF were found to regulate post-mating refractoriness, we next investigated whether these two neuropeptides functionally interacted. Single knockdown of *AT* or *sNPF*, as well as simultaneous knockdown of *AT* and *sNPF*, reduced the number of matings and extended the RS duration.

**Figure 5.**
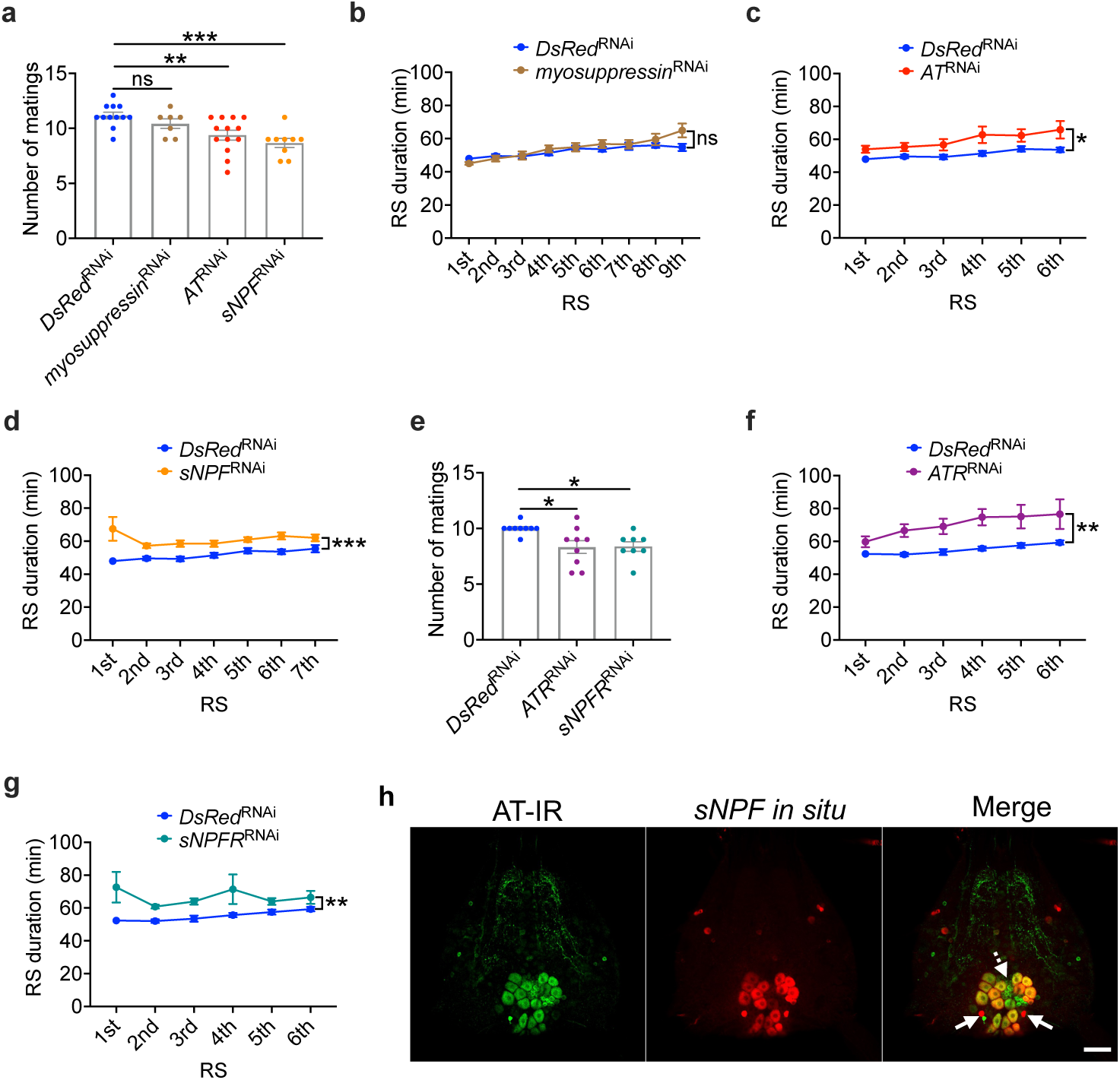
Regulation of male post-mating refractoriness by neuropeptide signalings. (a) Numbers of matings of RNAi male crickets injected with ds*DsRed* (*DsRed*^RNAi^), ds*myosuppressin* (*myosuppressin*^RNAi^), ds*AT* (*AT*^RNAi^), or ds*sNPF* (*sNPF*^RNAi^) in 12 h. Values are shown as mean ± SEM, n=7–13, Dunnett’s test, \*\**P*<0.01, \*\*\**P*<0.001. ns, not significant. (b-d) Duration of each RS of neuropeptide knockdown males. *Myosuppressin* (b), *AT* (c), and *sNPF* (d) knockdown males exhibited a minimum of 9, 6, and 7 RSs in 12 h, respectively. As a result, the durations of 9, 6, and 7 RSs are shown, respectively. Values are shown as mean ± SEM, n=7–13, two-way ANOVA, \**P*<0.05, \*\*\**P*<0.001. (e) Numbers of matings of RNAi male crickets injected with ds*DsRed* (*DsRed*^RNAi^), ds*ATR* (*ATR*^RNAi^), or ds*sNPFR* (*sNPFR*^RNAi^) in 12 h. Values are shown as mean ± SEM, n=8–9, Dunnett’s test, \**P*<0.05. (f, g) Duration of each RS of receptor knockdown males. Both *ATR* (f) and *sNPFR* (g) knockdown males experienced 6 RSs in 12 h at least. The durations of 6 RSs are shown in these two graphs. Values are shown as mean ± SEM, n=8–9, two-way ANOVA, \*\**P*<0.01. (H) Colocalization of AT peptide- and *sNPF* mRNA-positive cells in the male TAG using immunohistochemistry and *in situ* hybridization. Dashed arrows point to AT peptide-specific positive cell cluster. Solid arrows point to *sNPF* mRNA-specific positive cells. IR, immunoreactivity. Bar, 100 µm.

However, the simultaneous knockdown did not cause any additional or synergistic effects on the male post-mating refractoriness compared to single knockdowns (Supplementary Fig. 3).

Moreover, *AT* knockdown did not alter the transcriptional level of *sNPF* in the TAG, and *sNPF* knockdown had no effect on the transcriptional level of *AT* in the TAG (Supplementary Fig. 2b and 2c), suggesting that AT and sNPF might not be the downstream signaling molecules of each other. Therefore, it is likely that AT and sNPF parallelly regulate the post-mating refractoriness in male crickets through different targets of signaling pathways.

Most orthoptera species prepare a new spermatophore in the male reproductive system for the subsequent matings during the RS. In crickets, spermatophore materials, including seminal fluid from male accessory gland (MAG) and sperm from seminal vesicles (SVs), are transported to subgenital plate, where these materials are assembled into a mature capsule-like spermatophore^15^. Although previous studies suggest that the duration of the process of spermatophore preparation is a potential determinant of the duration of post-mating refractoriness in insects ^16,17^, it seems not to be the case in crickets. We removed MAG/SVs complex from the male adults on the first day after eclosion, preventing them from preparing spermatophores after sexual maturation. These MAG/SV-ablated males exhibited comparable RS duration to the control (Supplementary Fig. 4). This suggests that AT and sNPF regulate the refractoriness independently of spermatophore preparation, including the process of transport of seminal fluid and sperm from MAG/SV, in crickets. In *D. melanogaster*, silencing *Ecdysis-Triggering Hormone receptor* in the sole juvenile hormone (JH)-producing tissue, corpus allatum, reduces post-mating refractoriness in male flies, which can be partially restored by JH analog application ^18^. However, injection of a JH analog, Fenoxycarb, at various doses into crickets did not alter RS duration (Supplementary Fig. 5), suggesting the little effect of JH in the post-mating refractoriness in crickets.

We noticed that the RS duration increased after successively multiple matings (Fig. 5b and Supplementary Fig. 6a). The transcriptional responses of AT/ATR and sNPF/sNPFR signalings to multiple matings were examined. The transcriptional level of *AT* in the TAG decreased with multiple matings, while no alteration in *sNPF* expression in the TAG was observed (Supplementary Fig. 6b and 6c). However, changes in the secretion level of sNPF peptide cannot be ruled out. Since the physiological activities in the MAG and SVs were not required for maintaining the RS (Supplementary Fig. 4), AT and sNPF likely act on other tissues, such as the brain, which controls stridulatory behavior necessary for courtship ^19–21^. RNAi experiments confirmed *ATR* and *sNPFR* expression in cricket brains (Supplementary Fig. 2e and 2f). A slight increase in the transcriptional levels of *ATR* and *sNPFR* was observed in the brains of the multiply mated males compared to virgins (Supplementary Fig. 6d and 6f), indicating the potential involvement in feedback regulation.

We next examined whether AT and sNPF with similar effects on the male post-mating refractoriness were colocalized in the male TAG. Double-staining using immunohistochemistry for AT and *in situ* hybridization for *sNPF* were used. The specificity of AT antiserum was confirmed by dot blotting assay, preabsorption assay and colocalization of AT peptide and *AT* mRNA (Supplementary Fig. 7). AT peptide and *sNPF* mRNA were colocalized in the large cells with a diameter of around 25 µm in the posterior region and bottom region of the dorsal midline of the male TAG (Fig. 5h). These cells co-expressing AT and sNPF in the male TAG may be responsible for concomitant regulation of the post-mating refractoriness by AT and sNPF. Additionally, in the same regions, AT peptide was also specifically distributed in a group of smaller cells with a diameter of around 10 µm, while *sNPF* was specifically expressed in two other small cells with similar diameters (Fig. 5h). The different distributions of AT and sNPF in these small cells indicate that the two neuropeptides may have distinct functions, despite their concomitant regulation of the male refractoriness. Notably, the dorsal posterior midline cells expressing these neuropeptides in the male TAG seemed to be the male-specific dorsal unpaired median neurons directly innervating the reproductive tissues in crickets ^22^. Consequently, an arising question was whether AT and sNPF, expressed in different small cells in the male TAG, regulated the functions of distinct reproductive tissues.

### AT and sNPF target different sites within the male reproductive system

The male reproductive system in cricket encompasses testes, vasa deferentia, SVs, MAG, and ejaculatory duct (Fig. 6a). These reproductive tissues, innervated by TAG, play distinct roles in spermatophore preparation during the post-mating RS ^17^. To investigate whether AT and sNPF were involved in specific processes of spermatophore preparation, we explored the target tissues of AT and sNPF by examining the spatial distributions of *ATR* and *sNPFR* in the male reproductive system of the sexually mature male crickets. *ATR* exhibited the highest transcriptional level in the MAG, whereas the highest transcriptional level of *sNPFR* was observed in the vasa deferentia and SVs (Fig. 6b and 6c). The spatial difference in *ATR* and *sNPFR* expression in the male reproductive system indicated the specific regulatory roles of the two signalings in the process of spermatophore preparation. Considering the innervation of MAG and SVs by TAG, it was asked whether AT and sNPF functioned as neurotransmitters to regulate these reproductive tissues. Direct MALDI-TOF MS revealed that the mature peptides of AT and sNPF with theoretical *m/z* 1366.6 and 1331.8, respectively, were detected in the TAG-MAG/SV nerves (Supplementary Fig. 8), indicating their roles as neurotransmitters.

**Figure 6.**
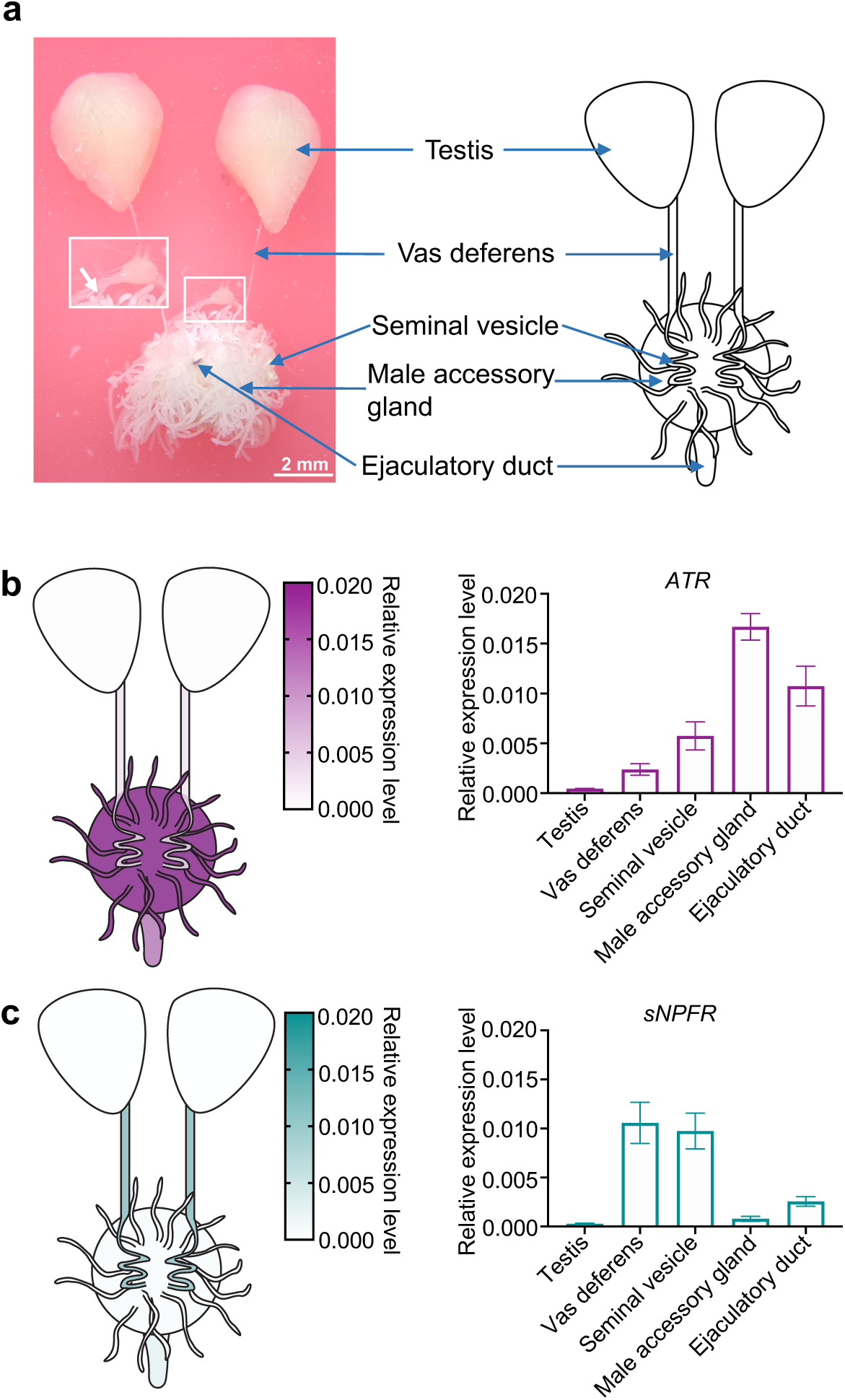
Different target sites of AT and sNPF within the male reproductive system. (a) Picture (left) and schematic diagram (right) of TAG-innervated male reproductive system in crickets. White box in the picture represents TAG innervation of male reproductive tissues which is enlarged. White arrow in the enlarged picture points toward the nerve projected from the TAG to the male accessory gland and seminal vesicle. (b, c) Left, the transcriptional levels of *ATR* (b) and *sNPFR* (c) in the male reproductive system indicated by heatmap. Right, the transcriptional levels of *ATR* (b) and *sNPFR* (c) examined by RT-qPCR. Values are shown as mean ± SEM, n=5.

The MAG of crickets emerged at the 3rd instar. From the 3rd instar to the early final instar, the MAG exhibited a morphology resembling a semitransparent faba bean. A significant morphological transformation was observed from the final instar to the adult stage. A substantial number of short or long, slender, blind-ending tubules protruded from the base of MAG (Supplementary Fig. 9a). Interestingly, the transcriptional levels of *ATR* in the MAG and *AT* in the TAG increased concomitantly with development. A notable surge was observed from the final instar to the adult stage (Supplementary Fig. 9a). This observation aligned with the period when the MAG underwent a drastic morphological change (Supplementary Fig. 9a). When the final instar and adult stages were further divided into 5 periods, synchronically gradual increases were observed in the size of the MAG and the transcriptional levels of *AT* in the TAG, as well as *ATR* in the MAG (Supplementary Fig. 9b). These results indicated the intimate association between AT signaling and MAG physiological activities. In contrast to the early emergence of MAG at the 3rd instar, the shaped SVs were absent until the third stage of the final instar (N3), when the terminals of vasa deferentia enlarged to form SVs. The SVs were empty at the final instar, and progressively filled with sperm after eclosion (Supplementary Fig. 9c). The transcriptional level of *sNPF* in the male TAG remained constant during the final instar but progressively increased after eclosion. The *sNPFR* mRNA level was higher in sexually immature adults than that in the final instar nymphs; however, the transcriptional level was not altered after eclosion, irrespective of sexual maturation (Supplementary Fig. 9c), suggesting the presumable function of sNPF signaling in SV physiological activities after eclosion such as sperm storage.

### AT signaling stimulates seminal fluid secretion from male accessory gland

As *ATR* and *sNPFR* were predominantly expressed in the MAG and SVs, respectively (Fig. 6), we delved into the specific roles of the two neuropeptide signalings in the physiological activities in these tissues. Initially, we examined the impact of silencing these two signalings on the physiological activity of the MAG. The weight of MAG in *AT* and *ATR* knockdown virgin males was comparable to that in the control virgin males (Fig. 7a), suggesting that AT signaling might not affect MAG growth. The MAG weight in the ds*DsRed*-injected males after successive mating for 24 h were much lower than that in the ds*DsRed*-injected virgin males (Fig. 7a and 7b). We observed that the MAG tubules in the virgin ds*DsRed*-injected crickets were filled with much milky seminal fluid, while the tubules in the mated ds*DsRed*-injected crickets were almost transparent (Fig. 7a and 7b), suggesting that multiple matings results in secretion of a great amount of seminal fluid for spermatophore formation, as observed in stalk-eyed flies, *Cyrtodiopsis dalmanni*^23^. The MAGs in *AT* knockdown crickets that underwent successive mating for 24 h were noticeably heavier than those in the control males (Fig. 7a). A similar effect was elicited by *ATR* knockdown (Fig. 7b). Considering the typical role of AT in the modulation of muscle contraction ^24^, the greater MAG weight in *AT* and *ATR* knockdown males after successive matings could be attributed to the inhibited MAG contraction, leading to the suppressed secretion of the MAG fluid. To verify this hypothesis, chemically synthesized AT was applied to the isolated MAG tubules immersed in Ringer’s solution. Exposure to AT stimulated the secretion of tubular contents in a dose-dependent manner (Fig. 7c and 7d). AT at final concentrations of 24 µM and 0.24 µM significantly increased the secretion (Fig. 7d), which was due to the noticeable contraction of the entire muscle epithelium of the MAG tubule (Supplementary Movie S1). Furthermore, application of cytochalasin D, an actin polymerization inhibitor, restrained the stimulatory effect of AT on the fluid secretion (Fig. 7d; Supplementary Movie 2), emphasizing the regulation of the MAG muscle contraction by AT. In contrast, RNAi targeting *sNPF* and *sNPFR* did not result in any observable changes in the MAG size regardless of the cricket conditions (Fig. 7a and 7b). Moreover, exposure to sNPF peptide did not induce fluid secretion from MAG tubules (Fig. 7c and 7d). Consequently, AT, rather than sNPF signaling, plays a crucial role in increasing seminal fluid secretion activity via stimulating muscle contraction of the MAG tubule.

**Figure 7.**
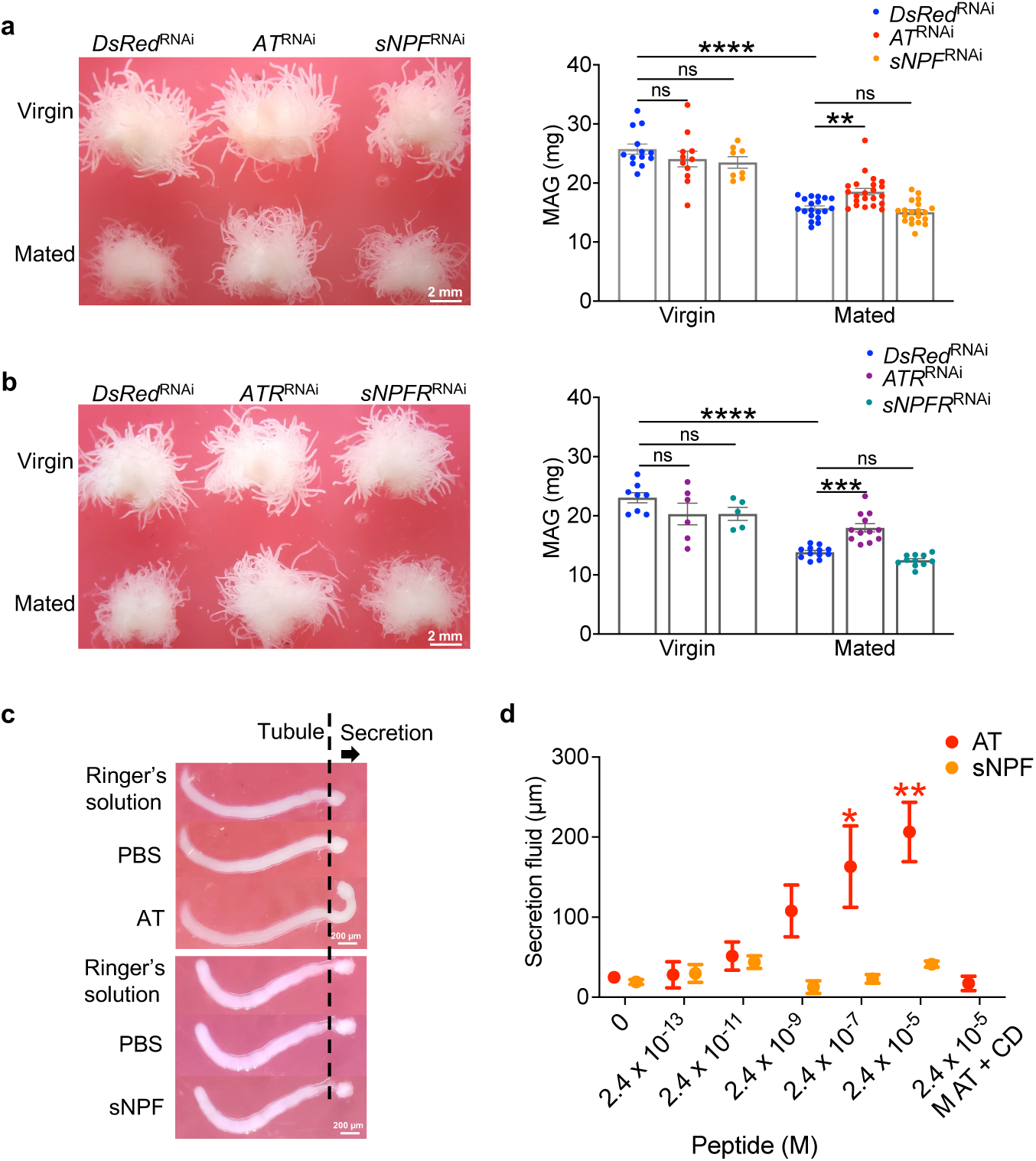
Regulation of MAG fluid secretion by neuropeptide signalings. (a, b) Weight of MAGs dissected from *AT* and *sNPF* knockdown (a), or *ATR* and *sNPFR* knockdown (b) virgin and successively mated males. Left, pictures of MAGs. Right, quantification of the weight of MAGs. Virgin; males with sexual maturity but without mating experience. Mated; males that finished successively multiple matings for 24 h. Values are shown as mean ± SEM, n=5–22, Sidak’s test for (a, b), \*\**P*<0.001, \*\*\**P*<0.001, \*\*\*\**P*<0.001. ns, not significant. (c) The pictures showing the isolated tubule with different treatments. (d) Effects of the synthetic AT and sNPF peptide application on the secretion of the fluid from the isolated MAG tubule and of cytochalasin D on the stimulatory effects of AT. An arrow points to the direction of fluid secretion. n=3, paired *t-*test, \**P*<0.05. ns, not significant.

### sNPF signaling promotes sperm storage in seminal vesicles

The predominant distribution of *sNPFR* transcripts within the male reproductive system was observed in the vasa deferentia and SVs (Fig. 6c), indicating that sNPF signaling may regulate sperm transport and storage. Like the seminal fluid in the MAG, the sperm in the SVs were also reduced by successively multiple matings (Fig. 8a and 8b). Sperm stored in the SVs was transported to form spermatophores. *sNPF* and *sNPFR* knockdown reduced the number of sperm in the SVs of both virgin and successively mated males (Fig. 8a and 8b). However, the number of sperm in spermatophores remained unaffected (Fig. 8c and 8d), indicating that sNPF signaling promotes sperm storage in the SVs but has no effect on the transport of sperm from SVs to spermatophores. The lack of influence in the number of sperm in SVs and spermatophores caused by *AT* and *ATR* knockdown suggests that the function of sNPF signaling differs from that of AT signaling (Fig. 8).

**Figure 8.**
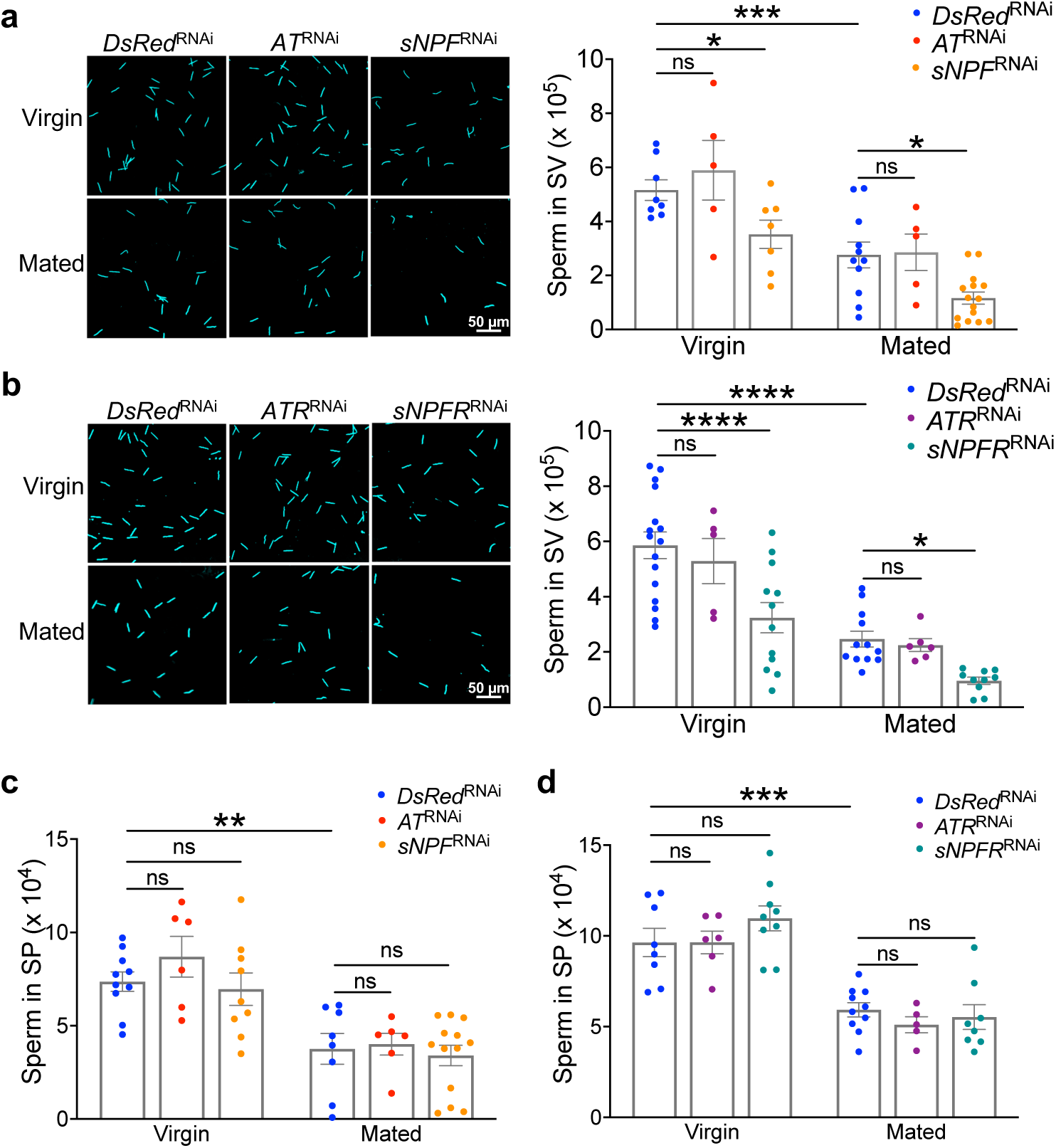
Regulation of sperm storage in SVs by neuropeptide signaling. (a, b) Effects of *AT* and *sNPF* knockdown (a), or *ATR* and *sNPFR* knockdown (b) on the number of sperm in SVs of virgin and successively mated males. Left, fluorescent pictures of sperm in seminal vesicles stained by DAPI. Right, quantification of sperm counts. (c, d) Effects of *AT* and *sNPF* knockdown (c), or *ATR* and *sNPFR* knockdown (d) on the number of sperm in spermatophores of virgin and successively mated males. Virgin; males with sexual maturity but without mating experience. Mated; males that have experienced successively multiple matings for 24 h. Values are shown as mean ± SEM, n=5-16, Sidak’s test, \**P*<0.05, \*\**P*<0.01, \*\*\**P*<0.001, \*\*\*\**P*<0.0001 ns, not significant. SV, seminal vesicle. SP, spermatophore.

Successively multiple matings decreased the MAG weight and sperm storage in the SVs (Fig. 7a, 7b, 8a, and 8b). Given that the physiological activities in the MAG and SVs were regulated by AT/ATR and sNPF/sNPFR signalings (Fig. 7 and 8), the effects of multiple matings on the expression of *ATR* and *sNFPR* in these reproductive tissues were also examined. However, the male crickets with multiple matings displayed comparable transcriptional levels of *ATR* in the MAG and *sNPFR* in the SVs to the virgin males (Supplementary Fig. 6e and 6g)

### sNPF signaling in males regulates post-mating behaviors of females

MAG and SVs in males may produce special materials transferred to females to regulate post-mating behaviors in some insect species ^25,26^. Considering the effects of AT and sNPF signalings on the MAG and SV functions, we evaluated the most typical post-mating behaviors, egg-laying and receptivity to remating, of females after mating with RNAi-treated males. The females mated with *AT* and *ATR* knockdown males laid comparable numbers of eggs with normal hatchability to the control and exhibited indistinguishable receptivity to another naïve virgin male (Fig. 9). In contrast, the post-mating behaviors of females were regulated by sNPF signaling in males.

**Figure 9.**
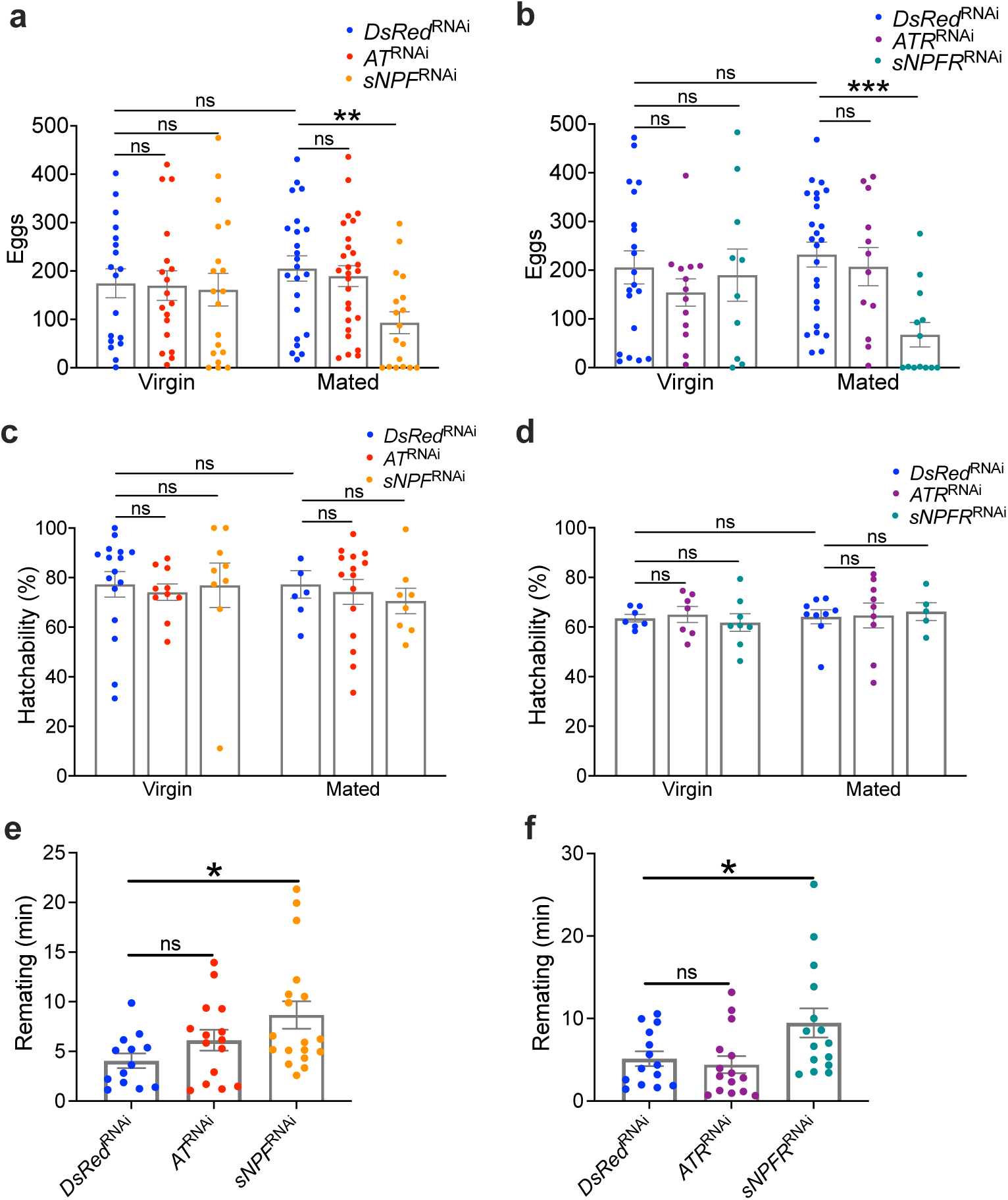
Regulation of post-mating behaviors of females by neuropeptide signalings in males. (a, b) Effects of *AT* and *sNPF* knockdown (a), or *ATR* and *sNPFR* knockdown (b) in males on the number of eggs laid by the females during 24 h after mating with males. (c, d) Effects of *AT* and *sNPF* knockdown males (c), or *ATR* and *sNPFR* knockdown (d) on the hatchability of the eggs laid by the females. Values are shown as mean ± SEM, n=5–27, Sidak’s test for (a–d), \*\**P*<0.01, \*\*\**P*<0.001. ns, not significant. SV, seminal vesicle. (e, f) Effects of *AT* and *sNPF* knockdown (e), or *ATR* and *sNPFR* knockdown (f) in males on the duration of remating of the females. Values are shown as mean ± SEM, n=13–18, Dunnett’s test, \**P*<0.05. ns, not significant.

Although the number of sperm transferred to females along with spermatophores were not affected by knocking down *sNPF* and *sNPFR* (Fig. 8c and 8d), the number of eggs laid by the females mated with *sNPF* and *sNPFR* knockdown males that had experienced successively multiple matings was significantly reduced (Fig. 9a and 9b), while the hatchability was not influenced (Fig. 9c and 9d). These results confirmed the little relevance between the number of sperm transferred to females along with spermatophores and the egg-laying behavior of females. Following successive mating with *sNPF* and *sNPFR* knockdown males for 24 h, females took longer to complete a successful mating with another virgin male, suggesting that their receptivity to remating was reduced (Fig. 9e and 9f).

## Discussion

AT and sNPF are pleiotropic neuropeptides in insects. Previous research on AT has predominantly focused on its regulatory roles in JH biosynthesis and muscle contraction ^27^; sNPF has been extensively studied for its involvement in feeding and metabolism ^28^. Several studies have hinted at the effects of AT and sNPF on oviposition in female insects ^29,30^. In contrast to these previous reports, our present study sheds light on the roles of AT and sNPF in the male-specific precisely timed post-mating refractoriness and the physiological activities in the male reproductive system, which have been little studied so far.

The present study revealed that *AT* and *sNPF* knockdown significantly extended the post-mating RS (Fig. 5), suggesting that the two neuropeptides play crucial roles in promoting mating behavior and terminating the refractoriness. Our findings indicate that AT and sNPF may independently control the post-mating refractoriness through distinct downstream pathways (Supplementary Fig. 3). This contrasts with the functional interaction between the two neuropeptides reported in the silkworm, *Bombyx mori*, where sNPF enhances feeding motivation by counteracting the inhibitory effects of AT ^31^. Furthermore, in *B. mori*, AT from the brain stimulates the expression of *sNPF* in the corpora cardiaca to suppress JH synthesis in the corpora allata, specifically at the early 5th instar ^32^. These reports and the present work indicate that the functional interaction of AT and sNPF differs according to the species, tissues, and developmental stages.

The process of spermatophore preparation in the male reproductive system appears dispensable for maintaining the post-mating refractoriness, suggesting the independent regulation of the refractoriness and spermatophore preparation in crickets. Such independent regulation of the concurrently occurring reproductive behavior and reproductive physiological activity by the same regulatory factor is also reported in *D. melanogaster.* Four abdominal ganglion interneurons expressing a neuropeptide, Corazonin, in male fruit fly independently control the copulation duration and the transfer of sperm and seminal fluid ^33^. If the male reproductive system for spermatophore preparation does not mediate the effects of AT and sNPF on the precisely timed refractoriness, there should be other target tissues of AT and sNPF by which the post-mating refractoriness is regulated. Brain is a potential target tissue since brain is the control center of stridulatory behavior in insects ^19–21^. A previous study suggests that TAG may secrete regulatory factors affecting the roles of brain in mating response during the RS in *G. bimaculatus* ^12^. It is speculated that AT and sNPF function as endocrinal regulatory factors that are secreted from the TAG to bind to their receptors in the brain to activate or inhibit downstream signaling, followed by initiation of courtship song and termination of the refractoriness. JH signaling can be excluded as the downstream of AT and sNPF in regulation of the refractoriness (Supplementary Fig. 5). Like crickets, male rat also exhibits post-mating refractoriness. Disruption of central serotonergic systems in the brain shortens the RS duration, whereas inhibition of central dopaminergic and noradrenergic systems increases the RS duration ^34–36^. Therefore, serotonin, dopamine, and octopamine, the monohydroxylic counterpart of noradrenaline in insects, presumably serve as the candidates of downstream molecules in the brain to mediate the effects of AT and sNPF signalings on the refractoriness in crickets, which remains to be elucidated.

The distinguishable distributions of *ATR* and *sNPFR* in the male reproductive system highlight the distinct roles of the two neuropeptide signalings in the physiological activities in the system. Silencing *AT* and *ATR* resulted in a greater MAG weight in crickets that have experienced multiple matings, attributed to the inhibited MAG tubule contraction that caused the less fluid secretion (Fig. 7), which is reminiscent of the myostimulatory function of AT in other insect species ^37–39^. The reduced mating frequency, resulting in less usage of MAG fluid in *AT/ATR* knockdown males, could not account for the greater MAG weight because the successively mated *sNPF/sNPFR* knockdown males with lower mating frequency possessed MAGs with the comparable weight with the control (Fig. 7a and 7b). sNPF signaling in males had no effect on the MAG but regulated sperm storage in the SVs and post-mating behaviors in females (Fig. 8 and 9). Although female crickets mated with *sNPF* and *sNPFR* knockdown multiply mated males received spermatophores with a normal number of sperm, they laid fewer eggs (Fig. 8c, 8d, 9a, and 9b). One explanation for the suppressed egg-laying behavior is the decreased viability of sperm in the spermatophore, probably caused by *sNPF* and *sNPFR* knockdown. Alternatively, SVs in males may produce a substance which functions like sex peptide, that is transferred to females together with sperm in *D. melanogaster* ^5^. This putative substance may regulate the post-mating behaviors of females, including egg-laying and receptivity to remating, and its production may be controlled by sNPF signaling in male crickets.

AT and sNPF together regulated the duration of the post-mating refractoriness but modulated different functions of the male reproductive system (Fig. 5, 7, and 8). The regulation of these male-specific behavior and physiological activities by AT and sNPF in *G. bimaculatus* may be attributed to their male-specific expression in the male TAG (Fig. 4), which precisely controls the RS, and innervates the male reproductive system ^12,22^. AT and sNPF were colocalized in the large cells in the dorsal midline of the male TAG, with additional specific expression in different small cells (Fig. 5h). This suggests that the large cells co-expressing AT and sNPF may regulate the post-mating refractoriness, while the smaller cells specifically expressing AT and sNPF may control the reproductive system; the group of small cells specifically expressing AT may regulate the seminal fluid secretion of MAG, whereas another two small cells specifically expressing sNPF may control the sperm storage of SVs. Future work is needed to prove this hypothesis.

In summary, this study unveils the neuropeptide regulation of the post-mating refractoriness and physiological activities in the reproductive system in male crickets. AT and sNPF signalings together regulate the duration of the post-mating refractoriness but influence different reproductive tissues. AT targets MAG, stimulating its seminal fluid secretion activity, while sNPF targets SVs, promoting sperm storage in SVs, and affects the post-mating behaviors of females.

### Methods Insects

*G. bimaculatus* were reared in plastic containers (55 × 39 × 32 cm) at 28 ± 1°C under a photoperiod of 16 h light and 8 h darkness. The diets were composed of rabbit food ORC4 (Orient Yeast, Tokyo, Japan) and cat food (Orient Yeast, Tokyo, Japan) at a weight ratio of 4:1.

### RNA extraction and reverse transcription-quantitative PCR (RT-qPCR)

Tissues from crickets were isolated in saline solution (0.9% NaCl in distilled water) under the microscope. Total RNA was extracted using TRI reagent^®^ (Molecular Research Center, Inc., Cincinnati, OH, USA) according to the manufacturer’s protocol. The total RNA was treated with RQ DNase I (Promega Co., Madison, WI, USA) to remove the genomic DNA, followed by purification using phenol/chloroform extraction and ethanol precipitation. The purified RNA was dissolved in 0.1% diethylpyrocarbonate (DEPC)-treated water to a concentration of 100 ng/µl. The resulting RNA was reverse-transcribed to cDNA using ReverTra Ace^®^ (Toyobo Co. Ltd., Osaka, Japan) for RT-qPCR. *G. bimaculatus Elongation factor* (*EF*) (GenBank accession number: ABG01881.1) was used as the reference gene. The sizes of the amplified fragments were 60-100 bp. The mixture of the template cDNA, THUNDERBIRD^®^ SYBR^®^ qPCR Mix (Toyobo Co. Ltd., Osaka, Japan) and the primers for RT-qPCR (Table S1) was reacted on a Thermal Cycler Dice Real Time System TP850 (Takara, Shiga, Japan) with the program: 30 s at 95°C, 40 cycles of 5 s at 95°C, and 30 s at 60°C; ended by a dissociation curve analysis at 95°C for 15 s, at 60°C for 30 s, and at 95°C for 15 s. For the amplification of *Kinin*, KOD SYBR^®^ qPCR Mix (Toyobo Co. Ltd., Osaka, Japan) was used due to its high GC contents in the sequence. The experimental evaluation of RT-qPCR was performed by confirming the resulting dissociation curves. The relative transcriptional levels were calculated using 2^−ΔCT^ method ^40^.

### Sampling from the crickets at the different growth and development stages

Generally, crickets experience 8 or 9 nymphal instars and an adult stage. The crickets at different growth stages were divided into 5 groups (1st-2nd instar nymphs, 3rd-6th instar nymphs, penultimate instar nymphs, final instar nymphs and adults) based on the size and stage-specific morphological appearances. The external genitals of females emerged at the 3rd instar. External wing pads emerged at the penultimate instar. The external genital and wing pads of the final instar nymphs became much longer than those of younger nymphs. Both sexes of the 1st and 2nd instar nymphs were mixed as one examining sample for RNA extraction because of their indistinguishable appearances (Supplementary Fig. 1).

### Whole-mount *in situ* hybridization

The primers for probe synthesis without T7 promoter (Table S1) were mixed with GoTaq® Master Mixes (Promega Co., Madison, WI, USA) and TAG-derived cDNA to amplify fragments of cDNAs encoding neuropeptide precursors with sizes of 300-600 bp by PCR. The PCR products were purified using QIAquick Gel Extraction Kit (Qiagen, Hilden, Germany). The purified DNAs were inserted into pGEM-T vectors (Promega Co., Madison, WI, USA) to construct cDNA-inserted, which were transformed into *Escherichia coli* to amplify. The forward primer without T7 promoter and the reverse primer with T7 promoter were used to amplify the target fragment from the cDNA-inserted plasmid. Then, the resulting PCR product served as the template, together with DIG RNA Labeling Mix (Roche, Mannheim, Germany) and T7 RNA Polymerase (Takara, Shiga, Japan) to synthesize antisense RNA probes. The forward primer with T7 promoter and the reverse primer without T7 promoter were used for the synthesis of sense RNA probes, the negative control. TAGs were isolated from male and female adult crickets about 1 week after eclosion and then fixed in 4% paraformaldehyde (PFA)/ phosphate-buffered saline (PBS) at 4°C overnight. After washing with PBS-0.1% Tween-20 (PBT), the TAGs were treated with proteinase K (5 µg/ml; Thermo Fisher Scientific, Waltham, MA, USA) for 10 min at room temperature and refixed using 0.2% glutaraldehyde in 4% PFA/PBS. The TAGs were next immersed in prehybridization buffer [50% formamide, 25% 20 x SSC, salmon sperm DNA (1 mg/ml; Thermo Fisher Scientific, Waltham, MA, USA), heparin (1 mg/ml), 0.1% Tween-20] at 50°C for 2 h, and then incubated in the hybridization solution (prehybridization buffer plus RNA probes at the final concentration of 1 ng/µl) at 50°C for 16 h. Next, the washed samples were incubated with the anti-DIG-alkaline phosphatase (AP)-conjugated antibody (Roche, Mannheim, Germany) at 4°C overnight. The samples were washed with staining buffer [Tris-HCl (100 mM, pH 8.8), NaCl (100 mM), MgCl_2_ (50 mM), 0.1% Tween-20], and then added to nitroblue tetrazolium and bromo-chloro-indolyl-phosphate solution (NBT-BCIP, 1:50 in staining buffer, Roche, Mannheim, Germany) to develop color in the dark. The signal was detected under a microscope.

### Chemical synthesis of neuropeptides

The amino acid sequences of mature AT and sNPF peptides are ‘GFKNVALSTARGF-NH_2_’ and ‘SNRSPSLRLRF-NH_2_’, respectively. The fluorenyl methoxycarbonyl (Fmoc) method was employed to synthesize the peptides as described previously with some modification ^41^. Rink Amide resin (Sigma-Aldrich, St. Louis, MO, USA) was used to provide an amide to the C-terminal of the peptides. Fmoc amino acids (Kokusan Chemical, Tokyo, Japan) were reacted with 1- hydroxybenzotriazole and *N,N’*-dicyclohexylcarbodiimide in *N*-methylpyrrolidone for the elongation of the polypeptide chains. Before the addition of a new amino acid residue, the Fmoc protecting group of the previous amino acid was removed by 20% piperidine in dimethylformamide. After the elongation, the resin and protecting groups were removed by treatment with a mixture of trifluoroacetic acid (TFA), dichloromethane, anisole, trimethylsilyl bromide, and 3,4-ethoxylene dioxythiophene with a volume ratio of 10:5:2:2:1 on ice for 1 h. The resulting peptides were mixed with diethyl ether and centrifugated for precipitation. The precipitants were dissolved in 0.1% TFA aqueous solution and desalted using a reversed-phase

Sep-Pak^®^ C18 cartridge (Waters Co., Milford, MA, USA) for fractionation by washing with 0.1% TFA, and then the peptide fraction was eluted with 60% acetonitrile in 0.1% TFA aqueous solution. The eluate was purified using a reversed-phase high-performance liquid chromatography (HPLC) (Jasco SC-802, PU-880, UV-875; Jasco Int., Tokyo, Japan) on a Senshu Pak PEGASIL-300 ODS column (4.6 mm i.d. × 250 mm; Senshu Kagaku, Tokyo, Japan) with a linear gradient of 0–60% acetonitrile in 0.1% TFA over 40 min at the flow rate of 1 ml/min. The chromatography was monitored by 225 nm. The purified synthetic AT and sNPF were confirmed by measurement with matrix-assisted laser desorption ionization time-of-flight mass spectrometry (MALDI-TOF MS) analyses using TOF/TOF 5800 System (AB SCIEX, Framingham, MA, USA). The measurement was carried out over the positive reflector mode with the 30 kV for the accelerating voltage. The mixture of the purified sample solution (1 µl) and the saturated solution (1 µl) of α-cyano-4-hydroxycinnamic acid (Sigma-Aldrich, St. Louis, MO, USA) in 60% acetonitrile/0.1% TFA was loaded on the target plate for the measurement of AT peptide (a theoretical *m/z* 1366.6) and sNPF peptide (a theoretical *m/z* 1331.8). The purified peptides were lyophilized and then dissolved in PBS.

### Antibody characterization

The polyclonal rabbit antiserum (Eurofins Genomics K. K., Tokyo, Japan) was generated against AT peptide with the sequence ‘NH2-C+GFKNVALSTARGF-CONH2’. Specificity of AT antiserum was tested by a dot-immunoblotting assay as previously described with a slight modification ^42,43^. PVDF membrane (Hybond-P; Amersham, Buckinghamshire, UK) was first rinsed in methanol and subsequently rinsed in water for hydration. One microliter of chemically synthetic AT and sNPF with different amounts dissolved in water was dotted on the PVDF membrane, followed by air-drying. Non-specific sites were blocked by soaking the membrane in Tris-Buffered Saline (pH 7.5)-0.1%Tween containing 5% skim milk (TBSTS) for 1 h. The membrane was then incubated with an AT antiserum (1:1000) in TBSTS for 1 h. After washing with TBST three times, the membrane was incubated with the horseradish peroxidase-conjugated secondary antibody (1:3000; Invitrogen, Waltham, MA, USA) in TBST for 1 h. A chemiluminescence reaction kit (ImmunoStar LD; Wako, Osaka, Japan) was used to develop the signals. The result revealed that the addition of AT, but not sNPF peptide, produced signals, indicating AT-antibody binding. The intensity of the spots decreased with reducing amounts of AT and concentrations of the antiserum (Supplementary Fig. 7a).

The antiserum specificity was further validated using a preadsorption assay ^44,45^. For immunostaining, TAGs were dissected and fixed in 4% PFA at 4°C overnight. The TAGs were then washed with PBS containing 0.5% TritonX-100 (PBSTr) and blocked at room temperature for 1 h using blocking buffer [10% goat serum (Invitrogen, Waltham, MA, USA), 2% BSA (Wako, Osaka, Japan), 0.5% TritonX-100]. AT was added to the blocking buffer containing the antiserum (1:1000) to achieve final concentrations of 73, 7.3, and 0.73 µM. The mixture was incubated at 4°C for 24 h, followed by centrifugation at 15000 rpm for 30 min. The supernatants, along with the untreated antiserum and preimmune serum, were used to incubate the TAGs at 4°C overnight.

After washing with PBSTr, the TAGs were incubated with the goat anti-rabbit IgG labeled with Alexa 488 (1:1000; Invitrogen, Waltham, MA, USA) at room temperature for 2 h, and then mounted with 50% glycerol/PBS. The signal was visualized using an FV3000 confocal laser scanning microscope (Olympus, Tokyo, Japan). The immunoactivity of AT was completely abolished by antiserum preadsorption with 73, 7.3, and 0.73 µM AT (Supplementary Fig. 7b)

Furthermore, the antiserum specificity was confirmed using double staining of AT peptide and *AT* mRNA. Whole-mount *in situ* hybridization and immunohistochemistry were performed to detect the colocalization. Following hybridization with RNA probe, the tissues were incubated with an anti-DIG-AP-conjugated antibody (1:1000) together with the rabbit antiserum against AT (1:1000) at 4°C overnight. The color of the signals for *in situ* hybridization was developed using Fast Red (Sigma-Aldrich, St. Louis, MO, USA) and checked under the microscope every half an hour to avoid overstaining. After washing with PBSTr, the tissues were incubated with the secondary antibody labeled with Alexa 488 at room temperature for 2 h. The signal detection was performed using a confocal microscope. The equivalent signals were obtained using immunohistochemistry for AT peptide and *in situ* hybridization for *AT* mRNA (Supplementary Fig. 7c).

### RNA interference (RNAi)

The knockdown-targeting cDNA fragments ranging from 300-600 bp were amplified by PCR using the cDNA-inserted plasmids. The primers used for PCR had a T7 sequence at the 5’ end, as listed in Table 1 S1. The double-stranded RNA (dsRNA) was synthesized using T7 RNA Polymerase (Takara, Shiga, Japan). The resulting dsRNA (10 µg) was injected into the abdomen of the final instar male crickets just after penultimate molting. All crickets finished final molting to become adults successfully, confirming that there was no developmental deficit caused by RNAi. Partial sequence encoding *Discosoma sp.* red fluorescent protein (*Ds*Red; GenBank accession number: AAT48428.1) was used as a control for RNAi experiments in this study. RNAi efficiency was determined by RT-qPCR on the fourth day after eclosion.

### Mating assays

Sexually matured crickets around 1 week after eclosion were used. To illuminate the effects of RNAi on RS duration, each dsRNA-injected male cricket was paired with 3 virgin females to reduce the influence of female refractoriness on the mating success by males during 12 h. The duration of RS was measured by observation on video-recorded crickets. The duration of total RS but not the separated RS1 and RS2 was measured because of the continuous physical and physiological stimulation from the accompanied females in the same container, ensuring the appropriately fixed RS1 and RS2 duration.

To study the post-mating behaviors of the females, each RNAi-treated male cricket was mated with a virgin female. After once mating, the mated females were reared individually in a plastic container with freely available food and water-soaked tissue for water supply as well as for oviposition. The number of eggs laid by females during the next 24 h was counted. The water-soaked tissue was placed in a new container. The number of hatched nymphs was counted for the calculation of hatchability activity. To investigate the receptivity to remating, each female that had been paired with an RNAi-treated male for 24 h was mated with a virgin male without any treatment. The duration of remating, from male courtship to successful spermatophore extrusion, was recorded to assess the receptivity.

### MAG secretion assay

MAG secretion assay was performed as previously described ^46^. MAGs were isolated from sexually matured virgin adult males about 1 week after eclosion. The tubule of MAG was cut at the base. The isolated tubule was immersed in 10 µl of Ringer’s solution [NaCl (140 mM), KCl (4.7 mM), CaCl_2_ (2 mM), MgCl_2_ (2 mM), HEPES-NaOH (5 mM, pH 7.1)]. First, PBS (10 µl) was applied to the tubule. After 1 min, the synthetic AT or sNPF in PBS (10 µl) was applied to the tubule. The length of secreted fluid was measured. The difference before and after addition of PBS was calculated as the control. The difference before and after addition of the synthetic neuropeptides was calculated as the secretion activity. To investigate the effects of cytochalasin D on the muscle contraction of MAG tubule, the isolated tubule was immersed in 100 µM cytochalasin D (Cayman Chemical, Ann Arbor, MI, USA) diluted in Ringer’s solution, followed by the application of PBS and neuropeptide.

### Sperm counts

Sperm were counted as described previously with some modifications ^47^. Seminal vesicles and spermatophores were isolated from virgin and successively mated males in saline solution and then transferred to PBS (1 ml and 100 µl, respectively). The sperm were released out of the tissues by sonication. One microliter of the sperm-suspended solution was loaded on a slide and air-dried. The sperm spot was then covered with 4% PFA/PBS and air-dried, followed by addition of 90% ethanol and air-drying. The sperm DNA was stained with DAPI (1 µg/ml; Wako, Osaka, Japan) in PBS at room temperature for 30 min. After washing with PBS, 50% glycerol/PBS were added to the slide which was then covered with a coverslip. The sperm were counted under the fluorescence microscope.

### Statistics and reproducibility

Data plotting and statistical analyses were performed using GraphPad Prism. Schematic diagrams were drawn using Adobe Illustrator. All the experiments in the figures were performed at least twice. The statistical analyses were described in the respective figure legends.

## Data availability

The source data behind the graphs in the paper are at following accession number (doi:10.5061/dryad.vhhmgqp1x).

## Acknowledgements

This work is partly supported by Grants-in-Aid for Scientific Research, KAKENHI, Grant Numbers 19H02967, 20K21304, and 21H02129.

## Author contributions

Z.Z. and S.N. designed this experimental project. Z.Z. performed all experiments, analyzed the data, and wrote the initial draft of the manuscript. S.N. supervised the study and co-wrote the paper. Z.Z. and S.N. discussed the results and reviewed the manuscript.

