## Supplemental Figure for "Two neuropeptide signalings regulate post-mating refusal behavior and reproductive system in male crickets"

**Supplementary figures**

**Supplementary Figure 1.** Development of crickets. (a) Pictures of male and female crickets at different growth stages. The ventral side of the segments in red boxes are enlarged. Red arrows point to female external genital. Inside the blue rectangles are wings. Bar, 2 mm. (b) Schematic diagrams of crickets at different growth stages. Wings are marked in blue. The female external genitals are marked in red.
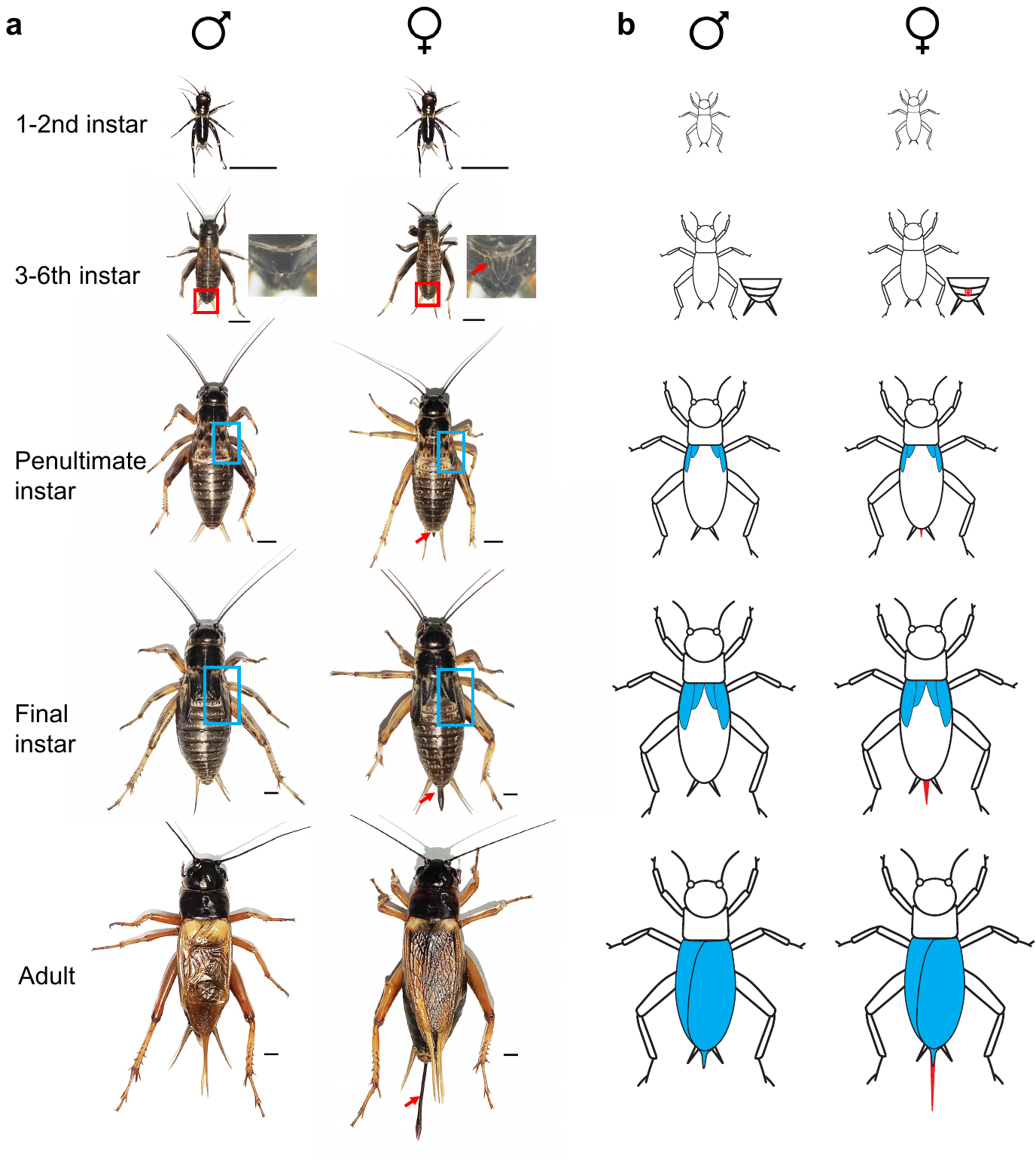

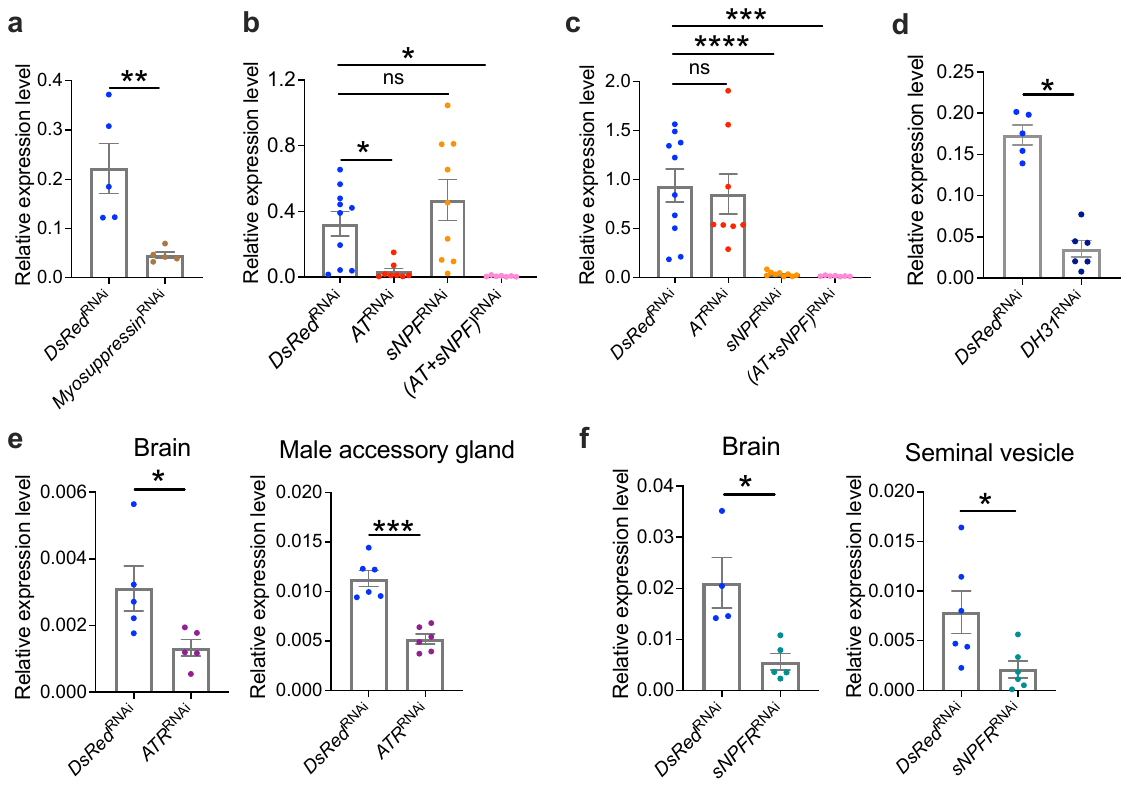
**Supplementary Figure 2.** RNAi efficiency. (a-c) Transcriptional levels of *myosuppressin* (a), *AT* (b), *sNPF* (c), and *DH31* (d) in the TAG. (e) Transcriptional levels of *ATR* in the brain and male accessory gland. (f) Transcriptional levels of *sNPFR* in the brain and seminal vesicle. Values are shown as mean ± SEM, n=4–10, unpaired *t*-test for (a), (d), (e), and (f), Dunnett’s test for (b) and (c), **P*<0.05, ***P*<0.01 ****P*<0.001, *****P*<0.0001. ns, not significant**.**

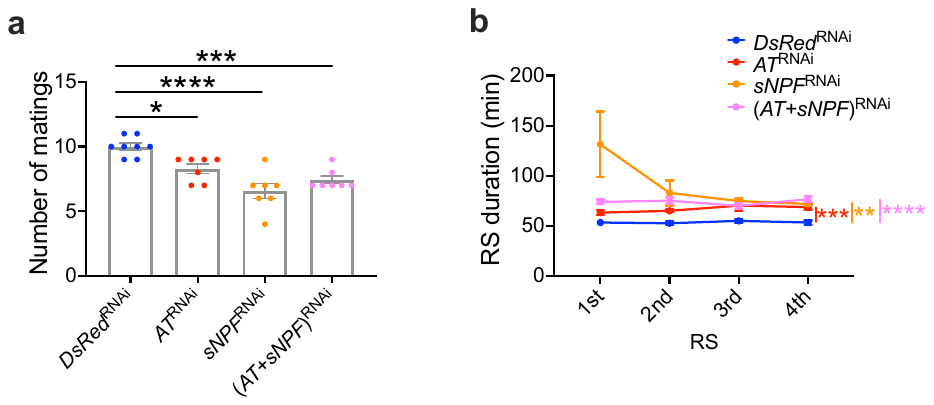
**Supplementary Figure 3**. Functional interaction between AT and sNPF. (a) Number of matings of males injected with ds*Dsred* (*DsRed*^RNAi^), ds*AT* (*AT*^RNAi^), ds*sNPF* (*sNPF*^RNAi^), or ds(*AT+sNPF*) [(*AT*+*sNPF*)^RNAi^] in 12 h. (b) Duration of each RS. Only the durations of the first 4 times of RSs are shown since all crickets experienced at least 4 RSs. Values are shown as mean ± SEM, n=7–8, Dunnett’s test for (a), two-way ANOVA for (b), **P*<0.05, ***P*<0.01, ****P*<0.001, *****P*<0.0001.

**Supplementary**
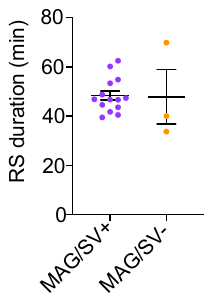
**Figure 4.** Effects of MAG and SVs on the duration of RS. MAG and SVs were removed from male crickets on the first day after eclosion. After one week, each male unable to prepare spermatophore was paired with one virgin female for 2 h. Those males completing copulation successfully entered a post-mating RS. The duration of the RS in MAG and SV-ablated males and that in the sham-operated males were compared. Values are shown as mean ± SEM, n=3–14, unpaired *t*-test.

**Supplementary**
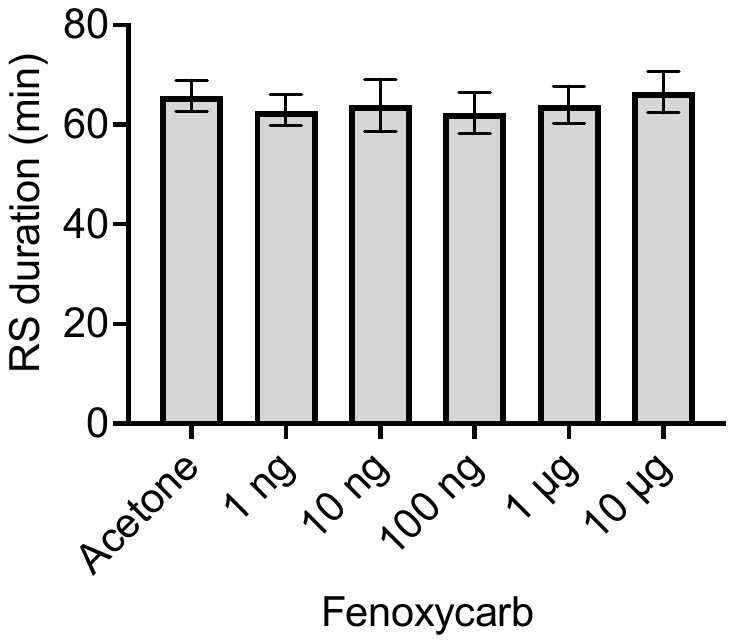
**Figure 5**. Effects of Fenoxycarb on RS duration. Values are shown as mean ± SEM. N=6-9, Dunnett’s test.

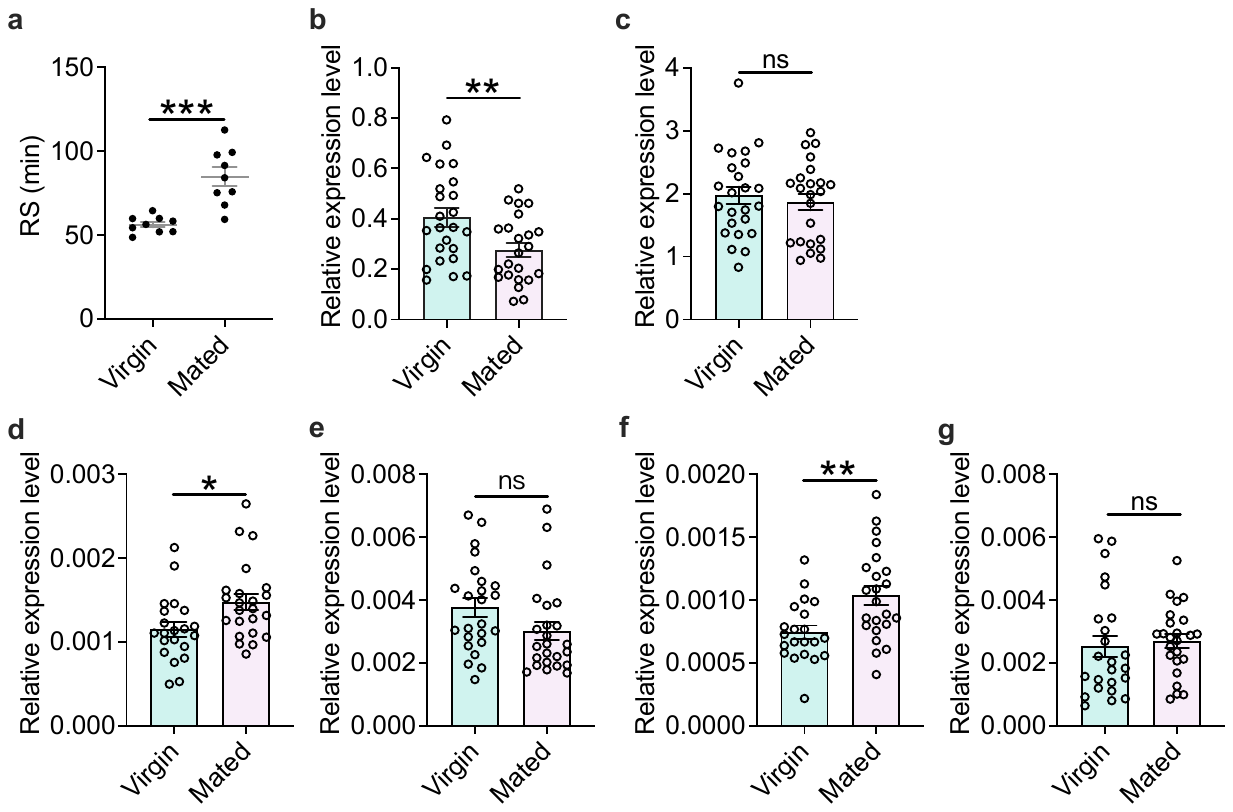
Supplementary Figure 6. Effects of multiple matings on the RS duration and expression of *AT*, *sNPF*, and their receptors. (a) Effects of multiple matings on RS duration. (b, c) Transcriptional levels of *AT* (b) and *sNPF* (c) in the TAG. (d, e) Transcriptional levels of *ATR* in the brain (d) and MAG (e). (f, g) Transcriptional levels of *sNPFR* in the brain (f) and SVs (g). Values are shown as mean ± SEM, n=21-24, unpaired *t* test. **P*<0.05, ***P*<0.01, ****P*<0.001. ns, not significant. Virgin; males with sexual maturity but without mating experience. Mated; males that finished successively multiple matings for 24 h.

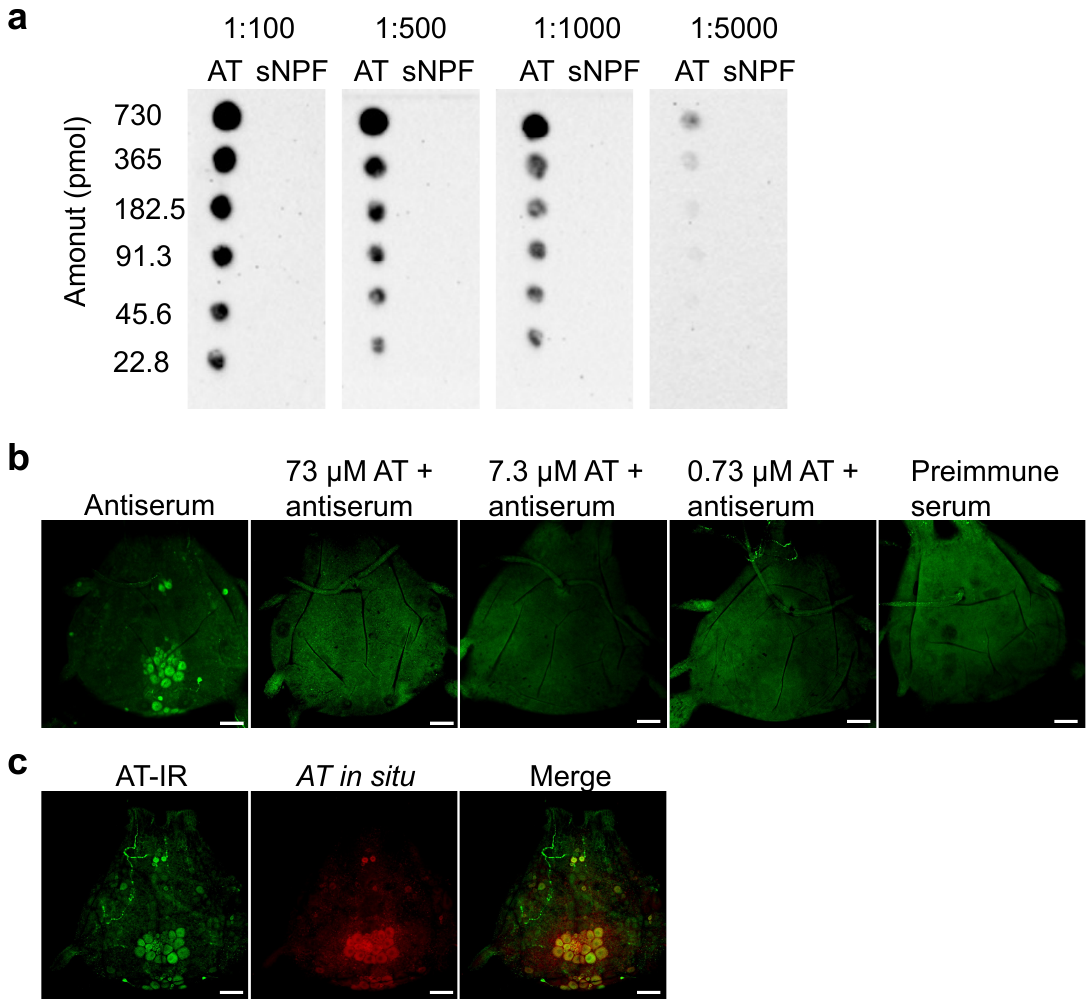
**Supplementary Figure 7.** Antibody evaluation and charaterization. (a) Dot blot assay. Synthetic AT and sNPF peptides with different amounts were used to react with the antiserum at different dilutions (1:100; 1:500; 1:1000; 1:5000). (b) Preabsorption assay. Synthetic AT with different concentrations were used to preabsorb the antiserum. Then, immunostaining of the TAGs was performed. (c) Double staining of AT peptide and *AT* mRNA using immunohistochemistry and *in situ* hybridization. IR, immunoreactivity. Bar, 100 µm.

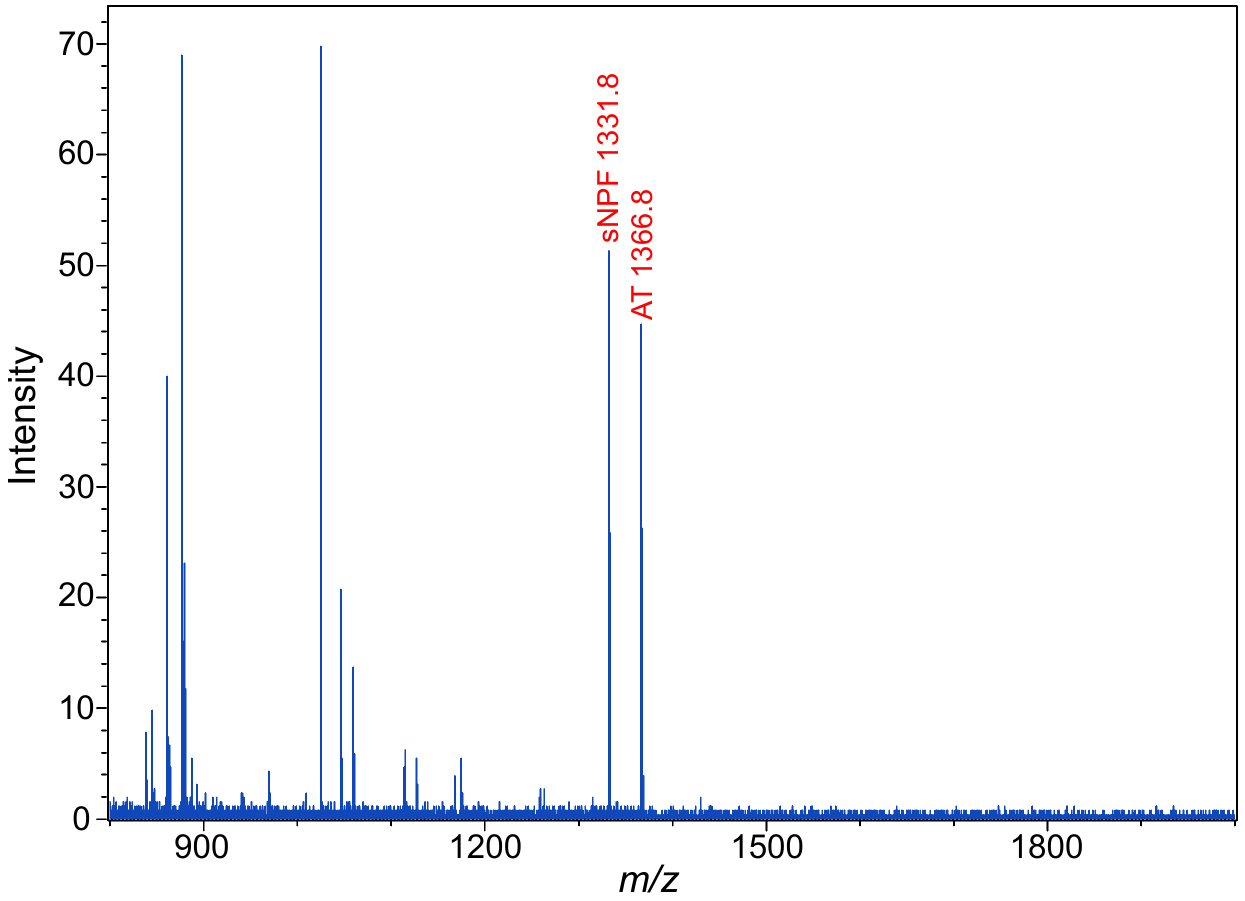
**Supplementary Figure 8**. Direct MALDI-TOF MS analyses confirm the existence of AT and sNPF mature peptides in the TAG-MAG/SV nerves.

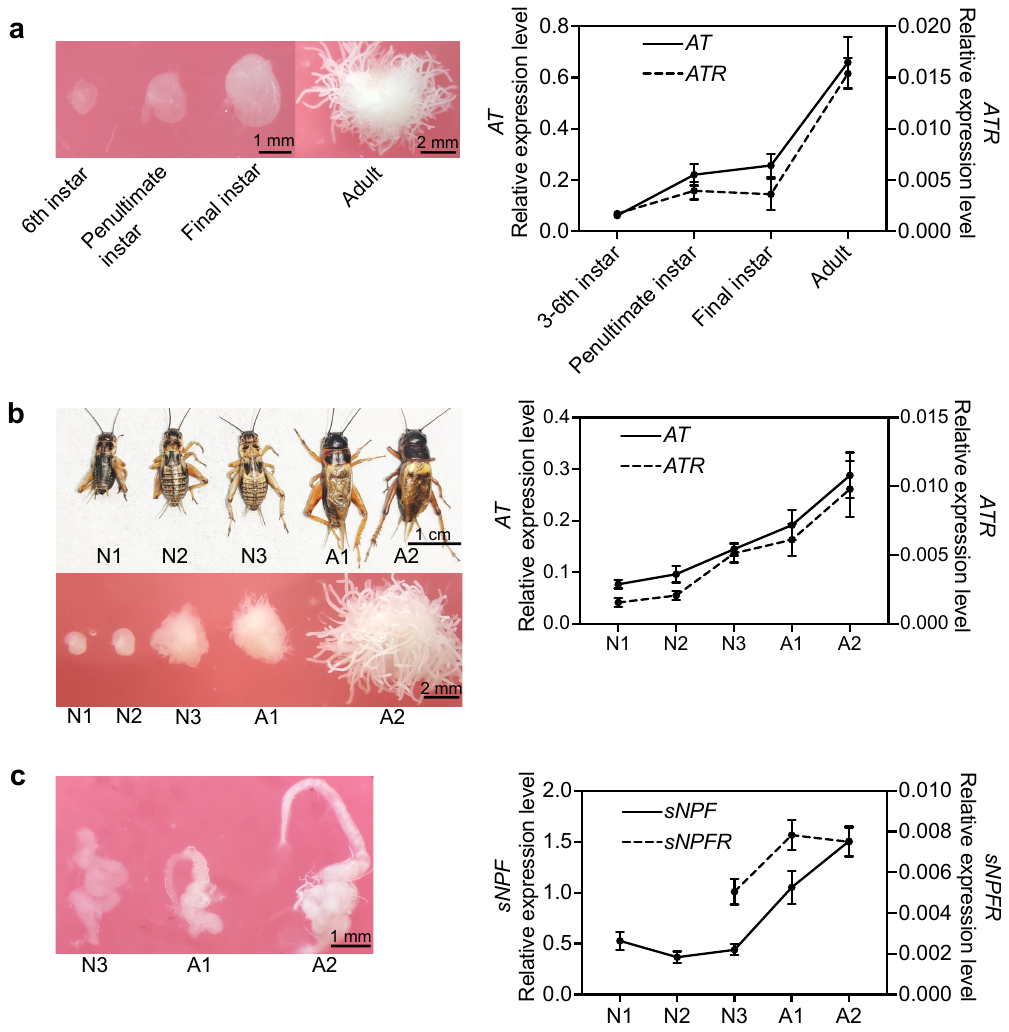
**Supplementary Figure 9.** Developmental changes in MAG and SV morphology and transcriptional levels of *AT*, *ATR*, *sNPF* and *sNPFR*. (a) Left, pictures of MAGs in crickets at different stages. Right, the transcriptional levels of *AT* in TAG and *ATR* in MAG with development. (b) Left, crickets in different periods of the final instar and adults (upper), and pictures of MAGs at the corresponding periods (lower). Right, the transcriptional levels of *AT* in TAG and *ATR* in MAG. (c) Left, SVs in crickets in different periods of the final instar and adult stage. Right, the transcriptional levels of *sNPF* in TAG and *sNPFR* in SV. The final instar and adult stage were further divided into 5 periods. N1, nymphs 1-2 days after penultimate molting with small and soft body; N2, nymphs 3-5 days after molting with a substantial largened body. N3, nymphs before eclosion with the wings separated from the abdomen. A1, adults 1-2 days after eclosion with soft body. A2, adults 1 week after eclosion. Values are shown as mean ± SEM, n=4-9.

**Supplementary Movie 1 (separate file).** Stimulatory effect of AT application on the MAG tubule contraction. The isolated MAG tubule was first immersed in 10 µl Ringer’s solution. Then 10 µl PBS was added. After 1 min, 10 µl AT peptide solution (1 µg) was applied to the tubule. The videos are played at 15x speed.

**Supplementary Movie 2 (separate file).** Inhibitory effect of cytochalasin D on the stimulated MAG tubule contraction by AT. The isolated tubule was first immersed in 10 µl Ringer’s solution with cytochalasin D at a concentration of 100 µM. Then 10 µl PBS was added. After 1 min, 10 µl AT peptide (1 µg) was applied to the tubule. The videos are played at 15x speed.

Table S1. Primers used in this study. F, forward. R, reverse. Probe, primers used for *in situ* hybridization. q, primers used for RT-qPCR. RNAi, primers used for RNA interference experiments.

| **Primer** | **Sequence 5’-3’** |
| --- | --- |
| AstA probe-F | AAGGCCGCATGTACTCCTTC |
| AstA probe-R | CACAGAATGATCGGCTCGGA |
| AstB probe-F | TTTCCGAGGCGTTTCGTCTT |
| AstB probe-R | TGCTTGCATTCGTAATCTGGT |
| AstCC probe-F | GACTACCAACGCTTCGACGA |
| AstCC probe-R | CACCGCCAATACACTCGACT |
| AstCCC probe-F | CTGGTGGACGACGATGGTAG |
| AstCCC probe-R | AAGAACGAGCTCGCACTGAA |
| Busicon a probe-F | CTCCTGTTCCTGGGCTCTTG |
| Bursicon a probe-R | CGGCATTCCAGAGGAGCTTT |
| Busicon b probe-F | CCCTTCCCAACAGCTCTCTG |
| Bursicon b probe-R | ACCCCAACTTCACTTGTGCA |
| CAPA probe-F | GCGATGCTCGTGTTCTTGTC |
| CAPA probe-R | TCCTGCATTGCTCTGAGACG |
| CCHa1 probe-F | ACCCCTTGCGTCTCTACAAA |
| CCHa1 probe-R | TATCAACAGGGCGAAAAAGG |
| CCHa2 probe-F | CAATGGCTGCAGAATTACCGG |
| CCHa2 probe-R | GTCCGTAACAACGCATGC |
| DH31 probe-F | TCTTGCAGTGCGATGGTGT |
| DH31 probe-R | CGCTTGTTCTCAACGTCGTT |
| DH44 probe-F | TCATGAACGAGCTCAACCGG |
| DH44 probe-R | GGTGACAGAACAGAGACGCA |
| FMRFa probe-F | TGCATCCACTCGACTTCCAA |
| FMRFa probe-R | GCTCACACAGGGGTTGAAGA |
| ITP probe-F | AGTAGGGCACGCACTTCAAA |
| ITP probe-R | AACGTGTAGAGCTTTGCCCA |
| Proctolin probe-F | AAAATTGCGGCTGGACCTTG |
| Proctolin probe-R | AATCTGAGACACTCCGACGC |
| myosuppressin probe-F | CGGCCTCTTCCTTCTCATCC |
| myosuppressin probe-R | GTGATCGACGTCTTGCCTCT |
| AT probe-F | CCATTCCCGGTCACTCAACA |
| AT probe-R | GCGCCCTCCTTATGTACACA |
| sNPF probe-F | GCAGAACTGAAAATGGCTGCT |
| sNPF probe-R | CGCTTGCGAACATACTCAGC |
| AstA q-F | GGGTCCCCATGTACGACTTC |
| AstA q-R | CACAGAATGATCGGCTCGGA |
| AstB q-F | GCTCTCCCGACCTTCTTGTT |
| AstB q-R | ACATCTTGAATCTTGACTTGGGAGA |
| AstCC q-F | TGTTGCAGCTCCGTCATGAT |
| AstCC q-R | GACTACCAACGCTTCGACGA |
| AstCCC q-F | TCAGTGCGAGCTCGTTCTTT |
| AstCCC q-R | TCTTTTAGGCCACGTGCGAT |
| Bursicon a q-F | TCCACTTTCTTGGCAGCACA |
| Bursicon a q-R | TCTTTTGCATGCACAGGACG |
| Bursicon b q-F | TGGAGTTCAGACGTGTGCTC |
| Bursicon b q-R | TAAAGTGGTGCCCGAAAGCT |
| CAPA q-F | AGGGTTGGACGATCAGGAGA |
| CAPA q-R | TCGTGGAAATGGGAAGAGGC |
| CCHa1 q-F | GTTTAGCCGAGAGAGGATGG |
| CCHa1 q-R | TTCACCATTCATGCAGAGGA |
| CCHa2 q-F | CAGCTGTGCAAGATGGCATC |
| CCHa2 q-R | CCGATTCCTGGGAGAAACGT |
| CCAP q-F | GTCAAACCATCGCTCGTGTG |
| CCAP q-R | CTTGCTGCAACAGAGGCAAA |
| Diapause hormone q-F | CGGAGAGACGTTCTGGTCG |
| Diapause hormone q-R | GTGAAACGCGATGGTGCTTC |
| DH31 q-F | AGATCTGCGACGTGGACATC |
| DH31 q-R | TTGAGGAAGATGCAGGGCTG |
| DH44 q-F | TCATGAACGAGCTCAACCGG |
| DH44 q-R | GTTCTGCTGGATGCGATTGC |
| FLRFa q-F | TCATAACAGGCATCGCGGTT |
| FLRFa q-R | AACAGCAATGTTTCGCGCAT |
| GPA2 q-F | CACAAGCGTTGATGCTCGAG |
| GPA2 q-R | ATAAGAACGTCGCACTGCCT |
| GPB5 q-F | CTTCGCTGGACTTGCAGACG |
| GPB5 q-R | GAGCGCTACGAGTTCATGCA |
| ILP q-F | GGGAGCTGCTCTTCAACAGA |
| ILP q-R | GAAAGAGCCAGAGCCCGAAT |
| ITP q-F | AGTAGGGCACGCACTTCAAA |
| ITP q-R | ACGTCCTCGAGACTCCTCTC |
| Kinin q-F | GGCCAAGCGCAACTTCAAG |
| Kinin q-R | GAGAAGTAGGCCTTGCGGG |
| NPF q-F | GTCCGCTGTCTCCTCATCTC |
| NPF q-R | GAGGAAGCCAGCAACGAGAA |
| Orcokinin q-F | CGATCGCACCGGATTCGATA |
| Orcokinin q-R | ATGAAGTTGTCGAAGCCGGT |
| Pyrokinin q-F | AAACCGTTGAGCCAAATCCG |
| Pyrokinin q-R | TCACAGTCGTCAACGTGCAA |
| Proctolin q-F | GGCGTACCTTCCAATTTGGC |
| Proctolin q-R | TTTGTCAACTGTCGACCGGT |
| Sulfakinin q-F | CTCGGCGAGATGAGCAAGAG |
| Sulfakinin q-R | GTGTCCGTAGTCGTCGAAGGG |
| ACP q-F | CCACCCATCTGCGGTAGAAG |
| ACP q-R | CCATGAGTTGCGGAGTAGCT |
| Mtosuppressin q-F | GCCTTCCCGGTCTCTCTCTA |
| Mtosuppressin q-R | GCGTGCGTGTGTCTTGAAAA |
| AT q-F | GATCGTGCACAAGTTCGTGG |
| AT q-R | GAGCAGTTCTTCCGGTGACA |
| sNPF q-F | GTCCCTGATGCTGCTACTCG |
| sNPF q-R | TCGTAGTCGCCGTAGGAGG |
| AT receptor q-F | CAGTAGGCGACGGACATGAA |
| AT receptor q-R | GCCGAGATGACCTTCACCAT |
| sNPF receptor q-F | GATCTTCGGCACCTTCACCA |
| sNPF receptor q-R | ACCTTCACGTAGCAGAAGGC |
| Elongation factor q-F | CCCTGCTGCTGTTGCTTT |
| Elongation factor q-R | CCCATTTTGTCGGAGTGC |
| myosuppressin RNAi-F | CGGCCTCTTCCTTCTCATCC |
| myosuppressin RNAi-R | GTGATCGACGTCTTGCCTCT |
| AT RNAi-F | CCATTCCCGGTCACTCAACA |
| AT RNAi-R | GCGCCCTCCTTATGTACACA |
| sNPF RNAi-F | CTCTTGCCACCAGGTTCCTC |
| sNPF RNAi-R | AGATGATCCCGTGAAGAGCG |
| AT receptor RNAi-F | ATGGTGAAGGTCATCTCGGC |
| AT receptor RNAi-R | GGCGGAGTACTACGACATCG |
| sNPF receptor RNAi-F | ACTCCGCTGTACTCCTTCCT |
| sNPF receptor RNAi-R | ATGAAGAAGGGCAGCACGAA |
| DsRed RNAi-F | AGAACGTCATCACCGAGTTCAT |
| DsRed RNAi-R | CCGATGAACTTCACCTTGTAGA |
| T7 promoter | GCTTCTAATACGACTCACTATAG |
